# Brain-wide monosynaptic single-neuron connectivity with ROInet-seq

**DOI:** 10.1101/2024.04.04.588058

**Authors:** Zhige Lin, Osnat Ophir, Thorsten Trimbuch, Bettina Brokowski, Muhammad Tibi, Hermann Schmidt, Christian Rosenmund, Hannah Hochgerner, Amit Zeisel

## Abstract

Viral projection tracing strategies help establish regional connectomes of mammalian brains. Monosynaptic connectivity tracing with G-deleted rabies virus (RV) establishes synaptic input connectivity, but cannot distinguish networks at cell resolution. We implemented a barcoded ΔG rabies virus to introduce unique molecular tags - and make network tracing amenable to readout by RNA-sequencing. First, we optimized and characterized library complexity and uniformity, such that detection of specific barcodes can reliably distinguish the individual monosynaptic input networks of multiple infected neurons in parallel. To deploy the method at scale; to hundreds of cells and full-brain volume per experiment, we developed regions-of-interest network sequencing (ROInet-seq); an accessible, scalable and low-cost spatial assay. ROInet-seq combines routine fluorescent imaging and processing of fixed tissue sections with a simple molecular biology workflow to capture barcode sequences in relevant regions-of-interest, and ultimately describes single-neuron networks brain-wide. In cortical brain areas the assay revealed preserved regional network motives, including co-inputs to single cortical neurons from distant and local sites. Towards improved spatial resolution and simultaneous detection of transcriptomes and networks we finally sampled barcoded ΔG rabies virus-infected hippocampus on commercial spatial transcriptomics assays and reveal details of the regions’ neurons local network architecture.

## Introduction

Neuronal networks are the biological basis for all higher functions and behaviours in animals. Tasks such as memory acquisition and retrieval, or sensing and processing social cues, require extraordinary flexibility, versatility and computational power. These capabilities are achieved by the brain’s multiregional circuits and local modifying networks of neurons, but revealing the architecture of their networks remains a major undertaking. Mapping neuronal networks has been approached across different scales of connectivity. At the most detailed level, great efforts are undertaken to reconstruct detailed maps of single neurons and their complete synaptic connectivity using transmission electron microscopy (Mahalingam et al. 2022; Turner et al. 2022; Yin et al. 2020), or expansion microscopy (Michalska et al. 2023; Tavakoli et al. 2025). Naturally, these approaches remain highly complex and costly. Combining electrophysiology with scRNA-seq (e.g., Patch-Seq, (Fuzik et al. 2016)) can measure neural connectivity and identity simultaneously, but is typically limited to several pairs of neurons. Long-range projections between distant brain regions cannot be recorded, making it challenging to obtain a complete map of single-cell neuronal connections. On the other end of the scale, viral projection tracing uses modified neurotrophic virus, such as fluorescent marker-expressing adeno-associated virus (AAV). Depending on the virus’ tropism, the region of interest’s input or output regions are then fluorescently labelled, revealing brain-region connectivity on a mesoscale level (Oh et al. 2014; Qiu et al. 2024). However, these projection maps generally do not reveal synaptic connectivity, and lack single-neuron resolution.

In sum, connectivity tools ideally map synaptic neuronal networks of single-neurons across substantial volumes of a mammalian brain; but typically navigate concessions between resolution and scale. Transsynaptic viral tracing strategies circumvent some limitations by visually labelling neurons that are synaptically connected to each other. To map synaptic input networks, a genetically modified virus (for example, G-deleted rabies virus (ΔG-RV) (Wickersham et al. 2007)), is injected to a site of interest (start site), and when provided with G *in trans*, can transfer to input neurons via its synapses. In this manner, ΔG-RV labels neurons infected in the start site -and their input neurons, revealing monosynaptic-connected input networks across full brain volumes *in vivo*. When multiple fluorophores and selective receptors are used, independent circuits can be revealed in mutliplex (Suzuki et al. 2020). Yet typically, multiple cells are labeled with the same fluorescence tag, which hinders distinguishing input networks of single neurons in the brain. Similar to a modification used in viral projection tracing (Chen et al. 2019; Kebschull et al. 2016; Qiu et al. 2024), several groups therefore recently developed a variant of ΔG-RV that carries a library of unique genetic barcode sequences. Barcodes are read by sequencing, and they increase the number of distinguishable markers that uniquely label individual networks in a single experiment. This allowed to track glial interactions (Clark et al. 2021) and monosynaptic retrograde neuronal connectivity *in vitro* (Saunders et al. 2022). *In vivo,* two groups (Zhang et al. 2023; Becalick et al. 2025) recently used *in situ*-sequencing to read network barcode sequences, and characterize the cells’ molecular identities, besides providing valuable spatial context of single cells in the network. This method is practically limited to small areas or local networks, and costly to implement at scale. Despite some limitations, the genetic barcoding strategy shows great potential in improving the resolution and scale of neuronal connectivity mapping.

Here, we generated high-complexity libraries of distinguishable rabies virus particles by inserting a 21-nt random barcode into the 3’UTR region of RV-*M* gene. We characterized barcoded ΔG rabies virus’ *in vivo* properties in droplet-based scRNA-seq and detected network barcodes and cell transcriptomes simultaneously We then implemented sequencing-based assays that navigate the concessions between resolution and scale, to resolve synaptic inputs to single cells. At the center of these efforts, we developed ROInet-seq; an accessible, coarse spatial assay compatible with standard histology- and molecular biology tools, to reconstruct single-neuron input networks at low sequencing costs. We also demonstrated how commercial spatial transcriptomics assays (Visium ST and STOmics Stereo-seq) can improve the method’s resolution of local networks in densely wired brain regions. Our work reveals quantitative and qualitative insights into cortical connectivity; including the robust occurrence of simultaneous inputs from multiple distal sites to a single cortical cell, and distinct, region-specific network cytoarchitectures.

## Results

To insert a 21-nt random barcode to the RV genome at an optimal position, we first tested the genome’s coverage in ΔG-RV-infected HEK cells using droplet-based 3’scRNA-seq by 10xGenomics. Of five genes in the ΔG-rabies genome (*N, P, M, GFP* and *L*), matrix protein-encoding *M* achieved the highest coverage, and we inserted the barcode at its 3’UTR (Fig. 1A). Then, rabies particles were recovered from HEK-cell transfection with the barcoded ΔG-RV plasmid library, and pseudotyped with the avian envelope EnvA (Fig. 1B, see Methods). For uniform representation of all barcodes in the libraries of RV plasmids and particles it was vital to eliminate bias introduced at any step of the protocol (Fig. 1C-D). Low-complexity and/or non-uniform libraries, such as when few barcodes are overrepresented, would lower the confidence that a given detected barcode in fact represents a single, initially infected starter cell. Therefore, we first introduced several measures to achieve highly upscaled, uniform RV plasmid and particle production (see Methods).

**Figure 1.**
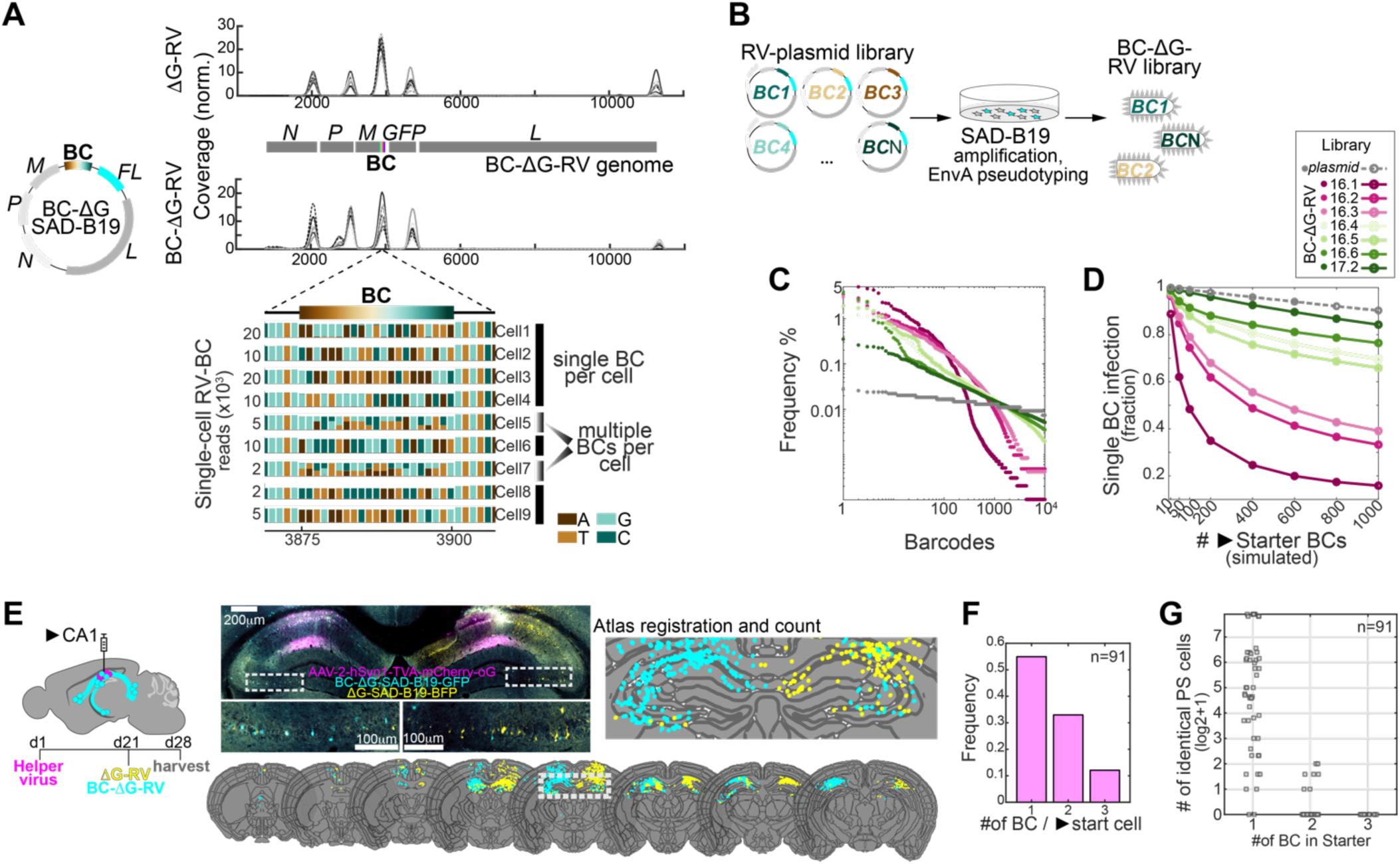
Generation of barcoded delta-G rabies virus and *in vivo* application for network tracing. (**A**). A 21nt random barcode was inserted to the 3’ site of the highly detected M-gene in G-deleted SAD B19 rabies virus. Right, coverage of the rabies genome in 10xChromium 3’ scRNA-seq in ΔG-RV, and BC-ΔG-RV. Below, reads in the BC region from 9 single BC-ΔG-RV-infected cells, detected by 10xChromium 3’ scRNA-seq. Most cells contained a single barcode, others contained multiple barcodes. (**B**) The BC-ΔG-RV library is recovered, amplified and pseudotyped with avian envelope (EnvA). (**C**) Frequency of the top 10^4^ detected barcodes in the RV-plasmid library and increasingly more complex BC-ΔG RV libraries. (**D**) Simulation of the expected cell fraction with singlet infection of increasingly more complex BC-ΔG RV libraries, and hypothetic comparison to the RV-plasmid, as a function of starter cells number. (**E**) Histological comparison of ΔG-RV and BCΔG-RV *in vivo*. Equal titers (0.8×10^8^IU/ml) of BC-ΔG-SAD-B19-GFP and ΔG-SAD-B19-BFP control, injected to unilateral CA1, each, display similar infection and transmission patterns. (**F**) In single-cell RNA-seq, frequency of detecting single or multiple BCs in a single starter cell (91 neurons). (**G**) In single-cell RNA-seq, scatterplot of 91 verified starter neurons, with 1, 2 or 3 unique BCs detected, and the indicated number of presynaptic cells with the same BC (combination).

For example, to maximize RV particles titer for *in vivo* tracing, plasmid-transfected HEK cells are typically amplified multiple times. However, during amplification, any fast-growing subset of cells would lead to an overrepresentation of the barcodes it produces. Highly amplified particles then occupy a large portion of the entire library (high frequency), and depending on sequencing depth, would in practice reduce overall library complexity (e.g., library 16.1, Fig. 1C-D). To overcome this limitation in subsequent RV-ΔG-BC libraries, HEK cells were transfected with the RV-plasmid library on a large, parallel scale and we limited production time to a single round, rather than amplifying cells multiple times (e.g., libraries 16.4, 16.5, 16.6, 17.2; Fig. 1C-D). This balanced cell growth, and drastically improved uniformity of the barcoded viral particles library. The optimized production strategy resulted in final libraries with acceptable uniformity: Measured by UMI-based amplicon sequencing, the top barcodes occupied ∼0.1-1% of the total library, and the probability of multiple cells infected by the same barcode at infection of around 200 starter cells was 10% or less (Fig. 1C-D).

We observed 2-6× lower titers of BC-ΔG-RV in virus production, hinting that insertion of a 21nt barcode to the 3’site of *M* may have compromised the virus’ replication or packaging capacity. Therefore, we next tested whether *in vivo* retrograde transfer efficacy may have been compromised, too. Indeed, compared to standard ΔG-RV, the barcoded ΔG-RV showed overall lower labeling efficacy of neurons in the mouse hippocampus CA1, and decreased efficiency of monosynaptic transmission to CA1 input cells (i.e., presynaptic-per-starter cells). At the same time, variability between animals and experimental batches was large for both virus variants (Fig. S1).

To assess how BC-ΔG-RV performs at cell resolution *in vivo*, we isolated BC-ΔG-RV-infected cells for single-cell RNA-seq (Fig. S2). We harvested one brain 8 days after BC-ΔG-RV injection to the CA1, then gently microdissected and dissociated fluorescent regions. Cell suspensions were quickly fixed, FACS sorted, and BC-ΔG-RV (GFP+) cells processed for scRNA-seq (10x Chromium). Out of 6310 recovered neurons, 66.4% cells contained BC-ΔG-RV transcripts, and 1.4% were confirmed starter cells (i.e., they also expressed genes derived from the AAV helper virus). This means that on average each starter cell transynaptically labelled 47.4 input cells (“presynaptic-per-starter”), which was in line with quantifications from imaging (Fig. S1). Starter cells were predominantly glutamatergic cortical-, or hippocampal pyramidal cells, although the lower transcriptome quality of RV-infected cells compromised finer subtyping (Fig. S2). We observed that in the start site, infection with multiple particles (multiple BCs) was nearly as common as single BC-infected starter cells (45% of all start cells had >1 BC, Fig. 1F, Fig. S2). And even that 34% of all RV+ cells had >1 BC (Fig. S2), most of these had BC combinations that did not overlap with those observed in any individual starter cell; but likely resulted from participating in multiple networks. Only in a minority of cases did we observe that multi-BC infected starter cells transferred more than one BC to their input networks (Fig. 1G). For example, out of 30 starter cells with 2 BCs, 76% transferred only one BC to input neurons. The same combination of 2 BCs was only detected in 7 input networks; and per network, in no more than 3 neurons; well below the average network size. Observing the same cell’s connection multiple times – due to multi-BC infection – thus likely accounts only for a minority of cases and is unlikely to overestimate connectivity. Finally, in the 91 sequenced starter cells we did not detect incidents of either multi-cell infection (i.e., one BC infects several starter cells) or connected starter cells (i.e., starter cell-to-starter cell transmission), as none of the 91 starter cells contained reads of the same BC sequence.

Next, we aimed to apply BC-ΔG-RV to characterize single-neuron input networks *in vivo.* Here, we assume that each barcode virtually represents a single neuron, based on the following observations: 1. Each start cell is dominantly infected by one or several barcoded RV-particles, where only one is likely to be transferred to most input neurons. 2. The BC library is highly complex, such that BCs can serve as a unique molecular tag. 3. BCs can be captured and reliably read by a universal RNA-seq protocol.

Therefore, with good characterization of BC-ΔG-RV *in vivo* performance and important improvements in library uniformity, we concluded that detection of a unique BC in a cell or region *in vivo* faithfully describes synaptic input of that cell or region to the start site. We argued that reconstructing input networks to a start site of interest required a sensitive, accessible method scalable to the full volume of the brain. Therefore, we developed “region of interest-network sequencing”, or ROInet-seq, where we specifically sequence all barcodes from individual BC-ΔG-RV-infected regions after full-volume fluorescent imaging. Briefly, 7d after BC-ΔG-RV surgeries, brains are harvested by transcardial PFA-perfusion, cryoprotected, cryosectioned (50μm), mounted and imaged (Fig. 2A-B). A semi-automated pipeline achieves full-brain registration to the Allen Mouse Brain Atlas and cell counting. Based on this analysis, ROIs are selected and microdissected (Fig. 2B-C). Each ROI is collected to lysis mix, where all transcripts are captured with an individually indexed poly-T capture oligo (ROI-index), reverse transcribed, and the RV-*M*-gene (flanked by the BC sequence) is specifically amplified (Fig. 2D-E, Fig. S3). Finally, ROIs are pooled and processed for sequencing (Fig. S3 and Methods). Each experiment results in an array of *M* counts (UMIs) of unique BCs detected in *N* ROIs. Further analysis can extract pairwise ROI BC overlaps, input networks of individual starter cells, and so forth.

**Figure 2.**
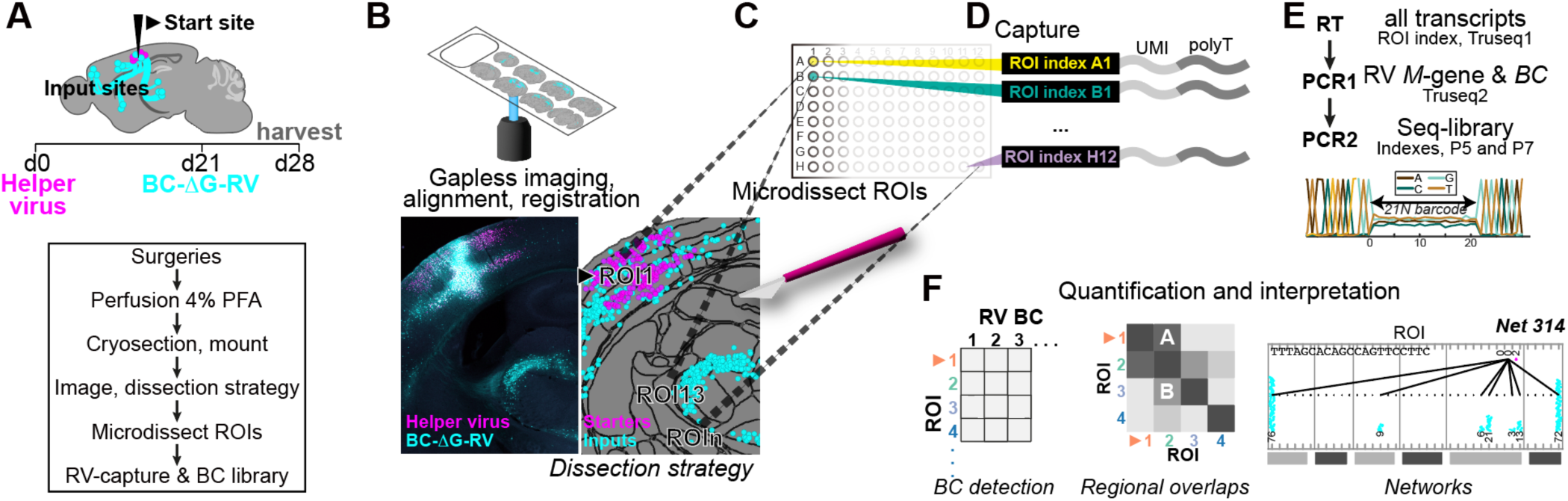
ROInet-seq for full-volume cell-resolved region-of-interest input network maps to the Somatosensory cortex (SSp). (**A**) Experimental outline; on top; helper virus (day 1), and BC-ΔG-RV (day 21) to the start site, and harvest day 28 by 4%PFA perfusion. Brains are postfixed, cryoprotected, cryosectioned, mounted; and imaged full-volume (**B**). All sections are aligned to the Allen Mouse Brain Atlas, and cells/ROInets are registered and quantified; informing the dissection strategy. Note; after registration (left sections), magenta represents starter cells (AAV+RV+). AAV+RV-cells are disregarded. (**C**) ROIs are microdissected and transferred to individual wells in a multiwell plate containing lysis buffer with barcoded polyT-capture oligos (**D**), to introduce a single barcode to all poly-adenylated of the same ROI. (**E**) Reverse transcription, targeted PCR to amplify the RV *M* gene with the BC fragment, and PCR for final sequencing libraries. Below, Pseudo-Sanger sequencing view of the nucleotide stretch containing the barcode sequence, as sampled by ROInet-seq. (**F**) Sequencing results in a list of unique detected ΔG-RV barcodes per ROI, with molecular counts, and allows reconstruction of (higher-order) monosynaptic input networks.

In three mice, we targeted the primary somatosensory cortex (SSp) (Fig. 3A). We imaged virus-labelled cells along the full brain volume and performed atlas registration and cell detection for all sections, resulting in per-region cell counts (Fig. 3B, Fig. S4). For all three brains, starter cells (AAV+RV+) primarily mapped to the deep layers of the barrel field (SSp-bfd), with some starter cells also detected in the adjacent nose field (SSp-n) (Fig. 3B). The majority of input cells (AAV-RV+) came from ipsi- and contralateral cortical sites within the SSp, the secondary somatosensory cortex (SSs) and the primary and secondary motor cortex areas (MOp, MOs), and ipsilateral subcortical regions as the thalamus (VP and PO) and striatum (CP).

**Figure 3.**
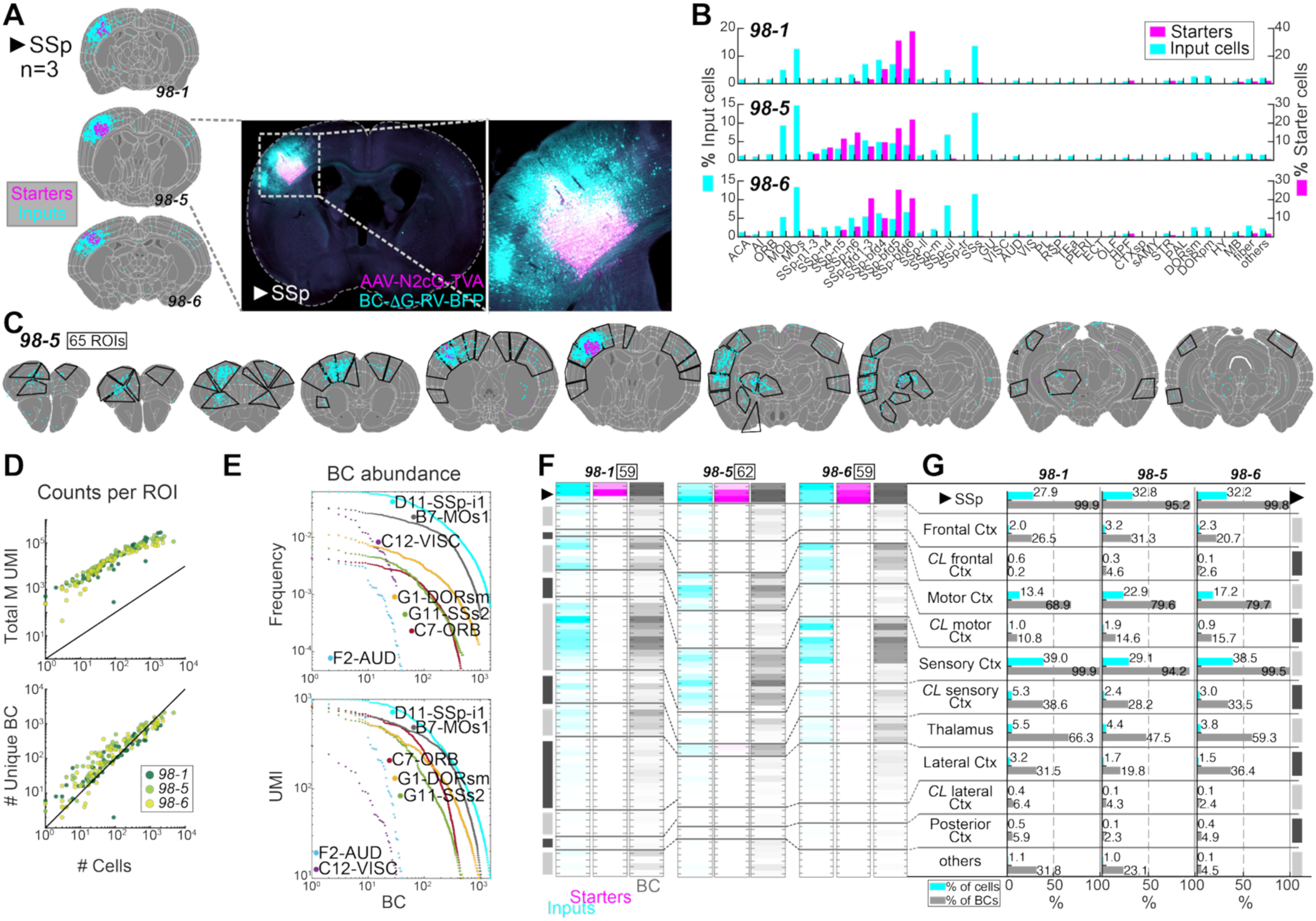
ROInet-seq for the primary somatosensory cortex (barrel field). (**A**) Monosynaptic input tracing to the mouse somatosensory cortex (SSp), in three mice (animal IDs 98-1, 98-5,98-6); showing coronal sections containing the SSp start site. Note; after atlas alignment and cell registration (left sections), magenta represents starter cells (AAV+RV+). (**B**) Quantification of starter and input cells after atlas registration, per experiment. (**C**) Example coronal sections of alignment and microdissection strategy (ROIs encircled), along the anterior-posterior axis. (**D**) For every ROI, counts of unique BCs and BC UMIs (sequencing), vs. automated cell counts (imaging). (**E**) BCs abundance frequency (as quantified by molecule counts, UMIs); shown for seven ROIs. (**F**) Per-ROI detection of input cells (cyan), starter cells (magenta) and BCs (grey; 98-1, 942; 98-5, 2764; 98-6, 1381), in the three experiments 98-1, 98-5, 98-6; normalized per column. ROIs are arranged and grouped by broad regional category; # of valid ROIs indicated next to experimental ID. (**G**) Percentage of cells (cyan) and barcodes (grey) detected in the full brain, per region top-category.

Based on the cell counts and atlas alignment, we designed a consistent microdissection strategy and collected >60 ROIs along the full anterior-posterior axis of the forebrain, per brain (Fig. 3C, Fig. S4, Table S1). All amplified barcode fragments were sequenced with unique molecular identifiers (UMIs). Size and network participation of ROIs greatly varied, but for ROIs in all three brains, the count of UMIs and unique barcodes both scaled with the number of cells detected per ROI (Fig. 3D, Fig. S5): On average, ROInet-seq detected ∼100 copies (UMIs) of 1 BC per cell; although this ratio varied between approximately 10-1000. We further observed that barcode detection was not binary. Instead – when considering individual barcode’s molecule counts – barcode abundances in all ROIs followed non-uniform distributions (Fig. 3E). This implied that RV-transmission was not uniform, and suggested that regarding barcode detection as a binary, rather than quantitative feature would be simplistic.

For each brain, we analyzed 941-2763 putative networks, based on the topmost widely detected unique barcodes (Fig. S4, Methods). Detection of BCs, starter and input cells was reproducible between ROIs of the three brains (Fig. 3F, Fig. S4). Unsurprisingly, almost all unique BCs were detected in the SSp start sites; but most unique BCs were detected in other ROIs, too (Fig. 3G, Fig. S4). This indicated that start cells usually received inputs from outside their immediate neighborhood (start site). Ipsilateral SSp ROIs were most connected, recapitulating observations of a recent RV retrograde tracing study, where single electroporated SS-bfd neurons received ∼80% of their inputs from local SSp cells(Inácio et al. 2025). Yet, other regions also showed great overlap: ROIs in motor areas contained between ∼70-80% of all unique BCs, thalamic areas ∼48-66% and contralateral somatosensory regions ∼30% of BCs. Outside of this “core” of SSp-input regions (Fig. 3F-G, Fig. S4), frontal cortex areas and lateral cortical association areas (including auditory, visceral, entorhinal) also each contained ∼20-30% of BCs (Fig. 3G, Fig. S4).

Compared to fluorescent cell counts alone, our per-ROI BC quantifications gave striking additional insights to SSp input network architecture. For example, outside the SSp start site, other ipsilateral somatosensory areas were most dense in input cells; making up ∼30% of all fluorescently labelled input cells (Fig. 3G). At the same time, these somatosensory areas also contained close to 100% of all BCs detected; meaning that the labeled neurons in these regions connected to almost all SSp starter neurons. In another example, thalamic regions together accounted for only ∼4.5% of fluorescently labeled input cells, but barcode detection suggested that these cells connected to ∼57% of all SSp start cells. Together, ROInet-seq uniquely revealed qualitative regional differences in input connectivity – that is missed in transsynaptic viral connectivity tracing with fluorescent labels alone.

Finally, to move from barcode detection to network mapping, we removed BCs at risk of multi-cell infection due to limits in library complexity and uniformity. We detected 941-2763 unique BCs in the SSp start site ROIs alone and therefore expected that >20% of BCs could infect more than one neuron (Fig. 1C-D, library 17.2). Since this would greatly hamper downstream interpretation of individual networks, we removed all BCs with frequencies >10^-4^ in either all three mice, or a deeply sequenced *in vitro* library (Fig. 4A). This step disproportionally removed BCs detected to label large networks more so than smaller networks (Fig. 4B), suggesting that more abundant BCs may have indeed infected multiple neurons in the start site that would then erroneously resemble large networks. The final pool of robustly detected, and unique BCs we analyzed labelled 628-2160 input networks that mostly spanned 5 or more ROIs, in all three experiments (Fig. 4C). In line with recent results describing brain-wide inputs to single, electroporated SSp barrel field neurons (Inácio et al. 2025), ROInet-seq thus uniquely revealed widespread patterns of multi-region inputs, to hundreds of single SSp starter neurons per experiment.

**Figure 4.**
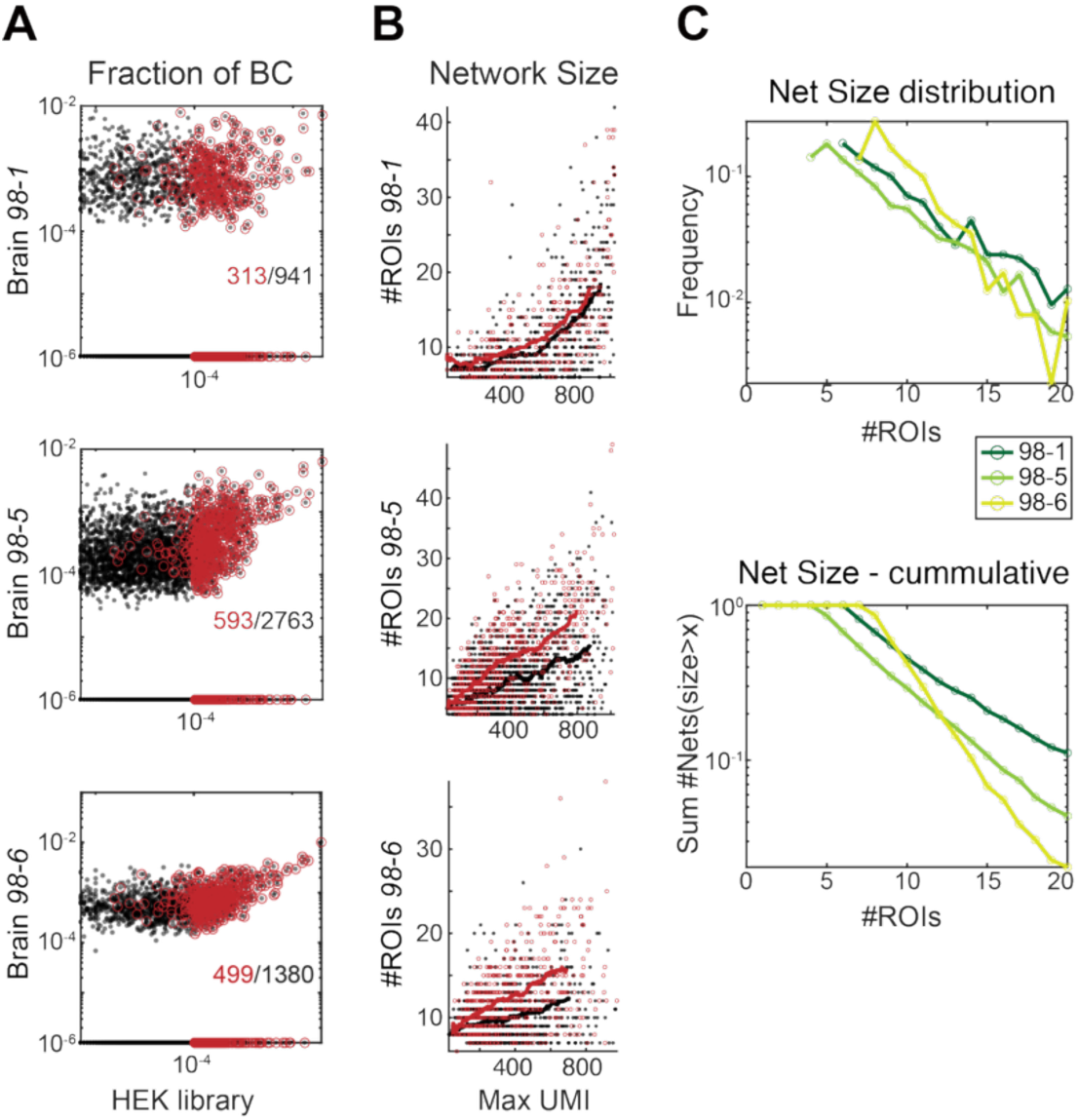
From ROInet-seq barcode detection to single-cell input networks. (**A**) Unique BCs detected at high abundances *in vivo* (animal IDs 98-1, 98-5,98-6) or *in vitro* were excluded for network analysis. Black, all BCs; red, abundant, discarded BCs. Numbers refer to BCs detected in the relevant experiment. (**B**) Network size distribution before and after removal of highly abundant BCs (red, as (**A**)); shown as quantification of unique BCs (x-axis) detected in # of ROIs (y-axis). Lines indicate BCs fit before (red) and after (black) removal. (**C**) Network size distribution after removal of high abundant BCs in the three animals as absolute (top) or cumulative (bottom).

We next examined individual neurons’ networks in detail. We plotted the detection of every individual BC in each ROI and visualized these SSp input networks together with BC molecule copy numbers (UMIs) (Fig. 5A, Fig. S6). Globally we observed that even that BCs described co-inputs from multiple ROIs, inputs from one region quantitatively dominated each network (Fig. 5B, Fig. S7). Most commonly, BCs were dominantly detected via somatosensory inputs and even locally in SSp start site ROIs. Dominant distal inputs came mainly from the motor cortex, and thalamus. These quantiative differences in distal input network architecture may reflect the starter neurons’ distinct functions: In retrograde RV tracing, single electroporated SSp-bfd neurons that were active in movement (whisker deflection) received dominant inputs from thalamus; while non-movement associated SSp-bfd neurons were more dominantly innervated by the motor cortex(Inácio et al. 2025).

**Figure 5.**
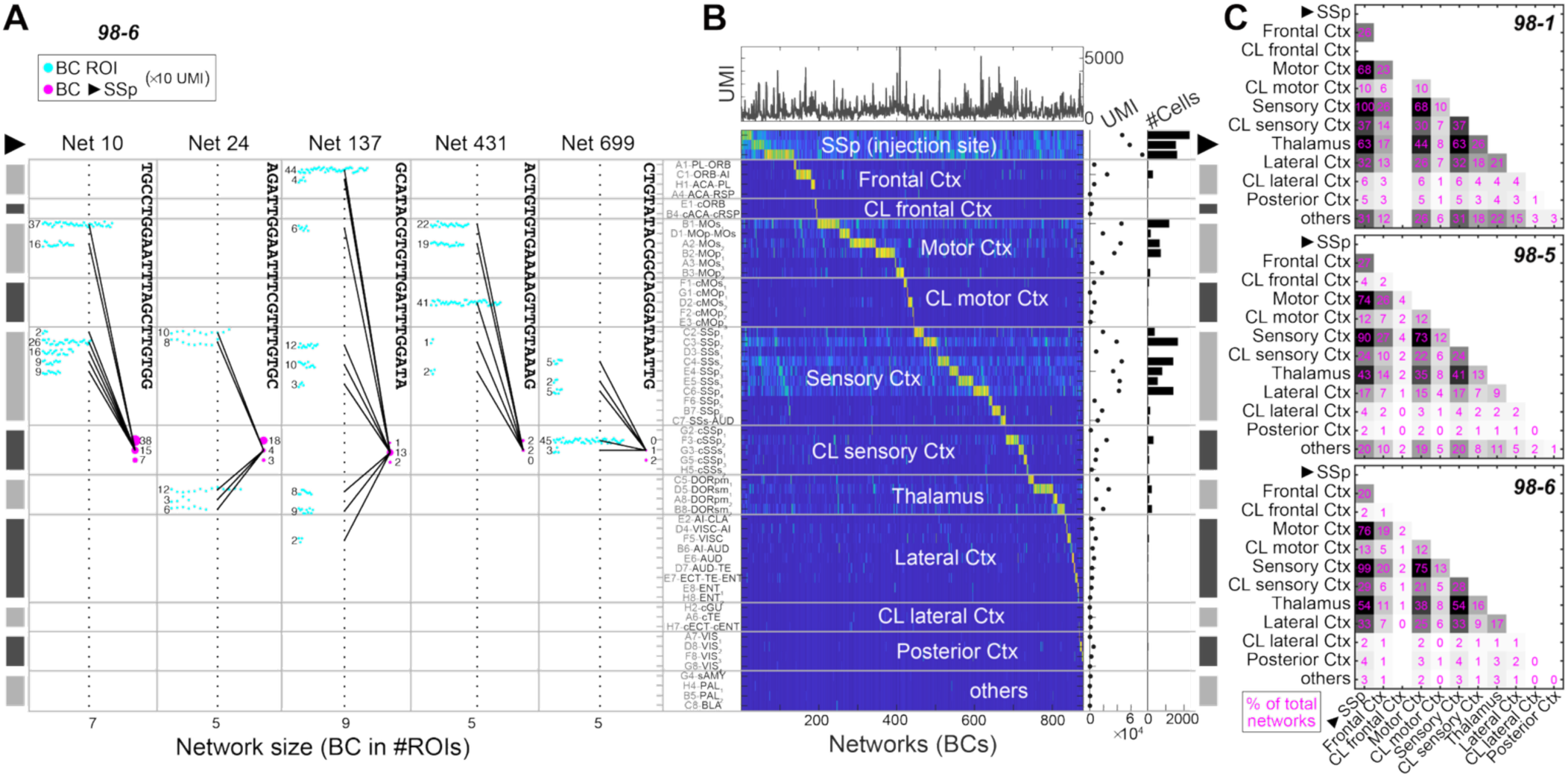
Quantification and convergence in SSp single neuron input networks. (**A**) Brain-wide input networks to five single SSp start site neurons in brain 98-6 at ROI resolution, visualized as dots, where each cyan dot represents detection of 10 unique network BC copies (UMI) in input ROIs, and magenta blob size represents unique BC copies in SSp start site. (**B**) Single-cell networks across 55 ROIs in brain 98-1, ordered by dominantly detected ROI. Colormap represents network BC copies (UMI) (blue, low; yellow, high). (**C**) Convergent regional inputs to single SSp start site neurons in brains 98-1, 98-5 and 98-6. Co-detection of network BCs between 12 regions (as annotated in (**B**)), in % overlap of all deteced network BCs. Greyscale represents minimum to maximum values standardized per ROI (excluding the diagonal).

We next examined pairwise shared BCs (Fig. 5C, Fig. S8) and found that several brain regions, and even functionally distinct areas often co-input to single SSp start cells. For example, over all experiments, 68-75% of BCs described co-inputs of somatosensory and motor regions, 41-63% were co-inputs of thalamus and somatosensory regions, and 35-44% were co-inputs from thalamus and motor cortex. The high frequency of these co-inputs may be unsurpsing, as these regions also gave most overall inputs to the SSp. However, compared to their frequencies, several other co-input motifs were overrepresented: For example, we found that rare inputs from contralateral cortex regions almost always overlapped with their respective ipsilateral regions: Contralateral motor cortex only consistutet only 10-13% of SSp inputs, and 10-12% of all SSp inputs came from both ipsi- and contralateral motor areas. Similarly, contralateral somatosensory regions accounted for 24-37% of all BCs, and the same fraction of all network BCs described co-inputs of ipsilateral and contralateral somatosensory cortex. Thus our analysis revealed overrepresented motifs of synaptic convergence of ipsi- and contralateral cortical regions with shared function, to single SSp neurons. Together, ROInet-seq uniquely detected and resolved higher-order input subnetworks of individual neurons from multiple remote sites; a connectivity property that is exclusively obtained only when single-cell resolution and brain-wide throughput are achieved in parallel.

To overcome the limited spatial resolution of ROInet-seq, we next applied BC-ΔG-RV for connectivity mapping on the commercial sequencing-based spatial transcriptomics platforms Visium ST (10x Genomics) (Fig. S9) and Sterero-seq (STOmics) (Fig. 6A). Given these platforms’ limits in throughput and scale we chose the hippocampus CA1 as start site, as most of its inputs come from within the hippocampal formation (CA1 and CA3). We injected BC-ΔG-RV in the CA1 (unilateral) and collected 10μm fresh-frozen cryosections to the capture areas. In Visium, a chip of four 6.5×6.5mm capture areas accommodated 16 coronal sections spanning the anterior-posterior axis of the hippocampus at 100μm spacing (Fig. S9). In Stereo-seq, three chips with a capture area of 10×10mm accommodated 27 coronal sections spanning the anterior-posterior axis of the hippocampus at 50μm spacing (Fig. 6A). For both platforms, we followed the standard protocol to generate transcriptome-wide cDNA libraries with spatial indices. For Visium, we additionally amplified the ORFs of AAV optimized glycoprotein (*oG*) and RV matrix (*M*)-gene (flanked in its 3’UTR by the BC sequence) (Fig. S9). The final sequencing library allowed us to detect transcriptome-wide gene expression, network BCs, and viral infection, spatially resolved to capture spots that in Visium contained several cells (diameter 55μm, center-to-center 110μm); and in Stereo-seq were of subcellular resolution (spot size ∼0.5μm). Neither platforms allowed for pre-assay imaging of native fluorescence, so viral infection was confirmed *post hoc*, using *in situ* hybridization (osmFISH) (Codeluppi et al. 2018) against RV M-gene on adjacent sections collected to separate slides (Fig. S9). In both assays, we detected elevated expression of inflammatory genes, such as *Cxcl13* and *Lcn2* (Fig. S9) in the section most focal to the injection site. In Visium ST, this section also featured an unexpectedly low number of BCs (Fig. S9); together hinting at tissue damage in the start site.

**Figure 6.**
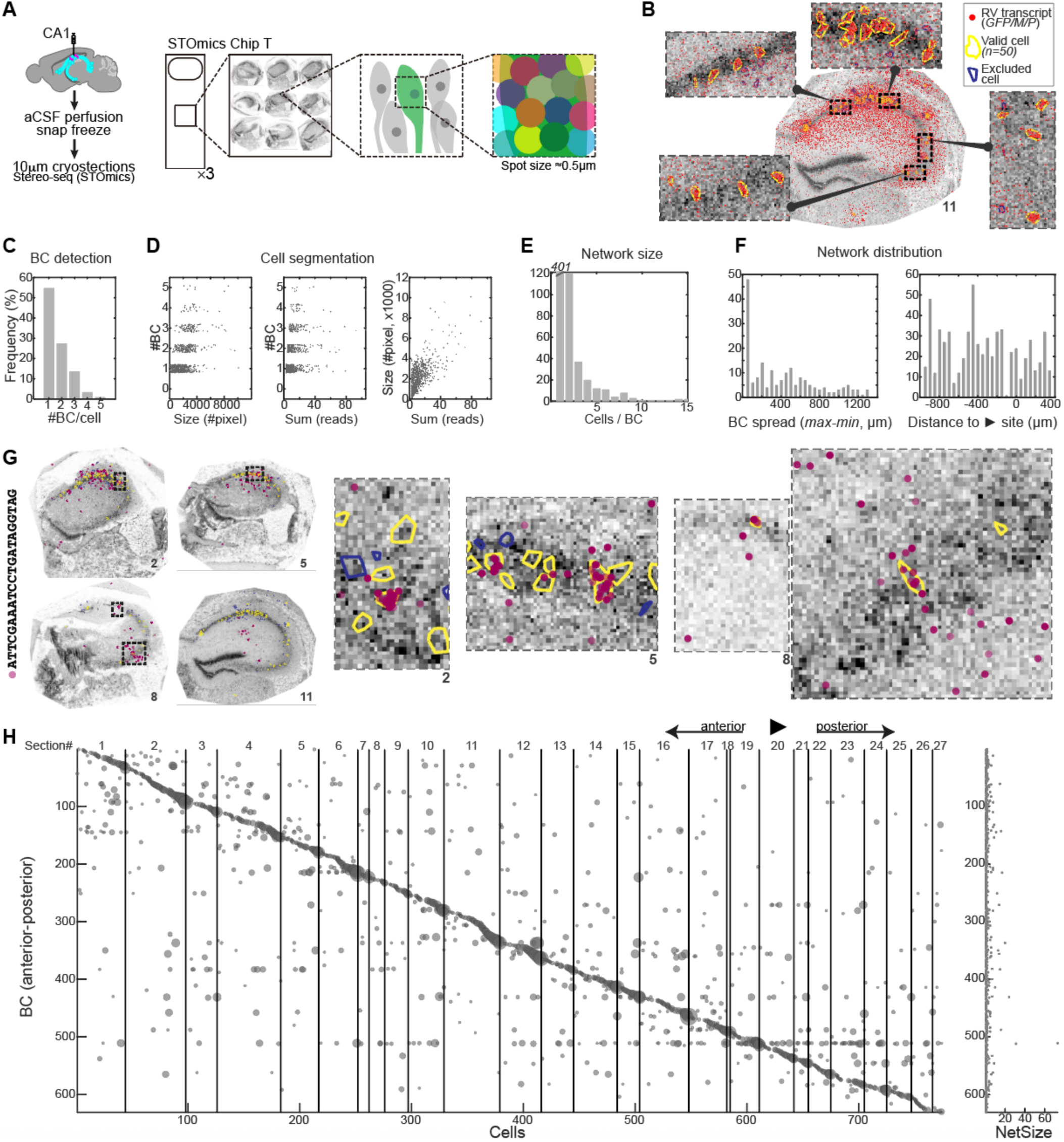
STOmics Stereoseq for hippocampus input network resolution. (**A**) Workflow for input network tracing in the hippocampus CA1 using the STOmics Stereoseq platform. We performed BC-ΔG-RV tracing after unilateral injection to the CA1, and collected 27 10μm fresh-frozen cryosections to 3 STOmics slides (10×10mm). (**B**) Network cell segmentation example in section 11 based on RV transcript clusters, detecting 50 putative single network cell soma (yellow), with zoom-ins on CA1 (top) and CA3 (bottom and right). Red dots, expression levels of RV transcripts *GFP, M and P* (bin 1); yellow outlines, segmented network cells; blue outlines, segmented cells below size or detection threshold. (**C**) Distribution of per-cell BC detection. (**D**) Number of BCs detected per segmented putative network cell as a function of cell size (left) and RV-reads (*GFP, M and P*) in the cell (middle); and network cell size as a function of RV-reads (right). Most segmented network cells contained a single BC; both BC detection and total RV-reads correlated with cell size. (**E**) Network size distribution, as number of cells per unique BC. (**F**) Anterior-posterior distribution of BCs in absolute distance (left) and relative to the start site (right). (**G**) Example of a single input network, visualized by BC detection (purple dots; BC sequence, left), in 4 sections. BCs detected outside the bounds of putative network cell somas (yellow outlines) are disregarded. Blue outlines depict segmented cells below size or detection threshold. (**H**) Anterior-posterior distribution and detection of individual BCs in all network cells, in section order. Blob size represents #BC reads.

In a total sampling area spanning 18177 capture spots, Visium ST detected 629 BCs distributed over 1848 spots. Most BCs were detected in 5 spots or less, and each BC spot commonly featured 1 or 2 BCs (Fig. S9). BCs were typically restricted within <200μm a.-p., although 13 BCs were detected across 1500μm sampled (Fig. S9-S10). The most abundantly detected BCs in the dataset, representing the best-connected starter neurons, typically also had the widest a.-p. spread, while lower detected BCs, representing less-connected starter neurons, often received inputs from a single site in the CA1 or CA3 that was typically anterior to the start site (Fig. S9-S10). These findings suggest that CA1 neurons can exhibit substantially distinct input network architectures; from single localized CA3 inputs, to remarkably widespread inter-CA1 inputs spanning large anterior-posterior distances (Fig. S10).

In one drawback, the spot resolution of Visium ST could not distinguish whether the detected BC spread stemmed from actual network cells, their projections (X-Y-Z), or even just diffusion of transcripts on the array (X-Y). And while we observed similar spread of RV transcripts on the Stereo-seq STOmics array, the assay’s subcellular spot resolution allowed for data-led cell segmentation: Reasonably-sized spatial spot clusters expressing RV-transcripts and BCs were interpreted as single labelled network cells (Fig. 6B, Methods) and we analyzed only BCs detected within the bounds of these likely cell bodies (Fig. 6B-C). Spurious detection of RV-transcripts (including barcodes) outside of spatial clusters was disregarded. Across 27 sections, we detected 774 network cells containing in total 631 unique BCs. In line with single-cell RNA-seq (Fig. 1F-G), most network cells contained a single BC (Fig. 6C). Cells with multiple BCs tended to be larger; reserving the possibility that some multi-BC cells may be unsegmented neighbor cells (Fig. 6B, D). Most BCs were detected in only one cell, but the method also detected 230 multicell (>1) networks, with 53 spanning 5 or more cells (Fig. 6E, Fig. S11). Networks were mostly restricted to ∼400μm in anterior-posterior distribution, with a slight bias to network labelling anterior to the start site (Fig. 6F); as exemplified also in Fig. 6G. RV genes were the most highly detected genes in network cells. In a subset of network cells, we also detected expression of transcripts that indicated their cell identity (Fig. S11). For example, glutamatergic markers labelled ∼142 of the network cells; and most expressed pyramidal marker *Neurod6*; with 124 positive for CA1-2 marker *Spink8*, and 38 expressing CA3 marker *Nptx1*, while only ∼20 cells expressed a general GABAergic signature (*Gad2, Gad1, Slc32a1*). More highly expressed neuropeptides *Sst* and *Npy,* marking GABAergic interneuron subpopulations, were better detected in ∼40 cells each.

Together, we demonstrated the application of single-cell input network tracing at spatial and single-cell resolution on two commercially available spatial transcriptomics platforms. In the hippocampus this revealed both locally restricted and wide-spread input network motives, and the platforms’ spatial resolution allowed to combine network information with cell location. In brain regions with complex and intermingled cellular diversity, these available methods already have the potential to reveal local input network architecture of molecularly defined cell types.

## Discussion

To systematically describe neuronal network architecture at single-cell resolution, we developed ROInet-seq, a scalable approach that combines monosynaptic input tracing with barcoded rabies virus. Each starter cell receives a unique barcode, and brain regions-of-interest that contain network cells (ROIs) are microdissected and sequenced in bulk. Because barcodes are unique per network, input networks to single cells can be reconstructed, at ROI resolution. ROInet-seq integrates seamlessly with standard histology, requires only simple molecular processing and shallow sequencing, and—using a highly complex, uniform viral library—resolves hundreds of input networks in parallel within a single experiment.

Applied to the somatosensory barrel field cortex (SSp-bfd), our results are coherent over independent experiments, and consistent with previous characterization of bulk input organization (Feldmeyer 2012; DeNardo et al. 2015). Beyond imaging-based quantification, sequencing revealed diverse connectivity patterns. For instance, although thalamic neurons accounted for only ∼4.5% of imaged presynaptic cells, they provided input to ∼57% of starter neurons, pointing at thalamic input divergence. Compared with *in situ* sequencing studies (Zhang et al. 2023; Becalick et al. 2025), ROInet-seq reconstructed larger brain-spanning networks and uncovered extensive convergence from multiple distant sites onto single SSp neurons. This included enriched motifs of simultaneous ipsi- and contralateral inputs from functionally overlapping cortical regions. Incorporating UMIs enabled molecular counting of barcode copies, providing quantitative measures such as regional input dominance. While variability in detection limits exact quantification, the linear scaling between UMI counts and imaged cells supports this interpretation.

Notably, several key findings were independently confirmed by Inácio et al. (Inácio et al. 2025), who combined two-photon calcium imaging in SSp-bfd with monosynaptic tracing from a single neuron per brain. First, the authors described expected massive local connectivity from within the SSp-bfd (∼80% of presynaptic cells); reflected in our findings via a large portion of BCs dominating within the SSp. Further, the authors described wide-spread, converging inputs from 6-7 larger brain regions; similar to our observed network sizes. Finally, Inácio et al. observed quantitative input dominance depending on neuronal functional state (i.e., movement-correlated, or not). It will be of great interest to test whether the quantitative input differences we observed in ROInet-seq may reflect similar functional specializations, and perhaps even network plasticity, of the traced postsynaptic neurons. These parallels suggest that ROInet-seq not only recapitulates quantitative and qualitative connectivity features but also extends them to hundreds of neurons simultaneously, providing robust and sensitive network maps at unmatched throughput. Importantly, the agreement with single-cell electroporation studies underscores the reliability of our approach.

We further applied ROInet-seq in combination with commercial spatial transcriptomics (Visium ST, STOmics Stereo-seq). In hippocampus, these assays enabled anterior–posterior sampling and higher spatial resolution of local BC-ΔG-RV networks. STOmics achieved single-cell resolution and confirmed *in vivo* infection parameters also observed in scRNA-seq, such as multi-barcode infection. However, both spatial transcriptomics and scRNA-seq suffered from sparse recovery of network cells—due to cell loss during dissociation, as reported previously (Zhang et al. 2023) or limited spatial scale—and reduced transcriptome quality of RV-infected cells. Consequently, detailed cell-type annotation of network members was limited. By contrast, ROInet-seq provided gapless, brain-wide coverage of inputs to SSp, albeit without capturing transcriptomes of individual presynaptic neurons.

Together, these findings establish BC-ΔG-RV tracing with ROInet-seq as an accessible, high-throughput method for single-cell resolved connectivity mapping, well-suited for comparative and functional neuroscience. Monosynaptic rabies tracing has already revealed circuit differences linked to sex dimorphism (Aloni et al. 2023), and behavioral circuit plasticity (Beier et al. 2017). The efficiency of monosynaptic rabies virus transfer was also shown not to depend on cell type or synaptic distance (Patiño et al. 2023). But monosynaptic input tracing likely underestimates overall connectivity, both due to inefficient retrograde synaptic transfer and cytotoxicity. Yet, recent advances – such as double-deletion B19 variants enabling long-term survival (Jin et al. 2024) and improved N2c substrains with higher efficiency and viability (Sumser et al. 2022; Reardon et al. 2016) – point toward further gains. Our use of N2c-glycoprotein helper virus yielded similar benefits, suggesting that continuing improvements in viral design will further enhance the sensitivity and functional relevance of BC-ΔG-RV.

### Limitations of the study

Like all approaches for reconstruction of synaptic networks, ROInet-seq involves several trade-offs. First, although UMI-based molecular counting provided valuable measures of regional input dominance, variability in detection prevents strictly quantitative estimates of connectivity strength. Second, the observed convergence and divergence patterns (e.g., thalamic inputs) cannot yet be disentangled from potential biases introduced by barcode detection or viral transfer efficiency. Third, the method sacrifices transcriptomic and refined spatial information of presynaptic neurons in exchange for brain-wide coverage and throughput, leaving the connectome–transcriptome link unresolved. Fourth, rabies-based tracing itself remains limited by incomplete retrograde transfer and cytotoxicity, which constrain the recovery and fidelity of network neurons. Finally, the bulk ROI-based design lacks the single-cell anatomical precision achievable with spatial transcriptomics or *in situ* methods, though these remain hampered by cost, tissue compatibility, and scale. Given the immense diversity of neuronal cells in the brain readily characterized by single-cell and spatial transcriptomics tools (Bakken et al. 2021; Siletti et al. 2023; Yao et al. 2021; Zeisel et al. 2015; 2018), the transcriptome-connectome link will be critical to implement.

Despite these limitations, our results demonstrate that BC-ΔG-RV with ROInet-seq provides robust, reproducible maps of neuronal input networks at unprecedented throughput. Future improvements in viral design, integration with transcriptomic profiling, and advances in spatial transcriptomics are likely to further close the gap between connectomic resolution, molecular identity, and functional interpretation— bringing brain-wide single-cell network reconstruction within reach.

## Acknowledgements

This research was supported by the European Research Council (TYPEWIRE-852786 to A.Z.) and Human Frontiers Science Program (CDA-0039/2019-C to A.Z.). We thank the FACS unit of the Technion LS&E Infrastructure Center for technical assistance, and former Zeisel lab member Etay Aloni for critical discussion.

## Author Contributions

L.Z., H.H. and A.Z. designed the study and planned experiments. L.Z. performed *in vitro* validation, viral injections, histology and ROI-microdissections, osmFISH, Visium ST and Stereo-seq. O.O. performed ROInet-seq, *in vitro* validation, Visium ST and Stereo-seq. T.T., B.B. and C.T. designed and performed RV library preparations and optimizations, H.S. constructed and produced RV plasmid libraries. H.H. performed scRNA-seq. L.Z. and A.Z. analyzed *in vitro* validation data and ROInet-seq data, A.Z. and H.H. analyzed scRNA-seq data, M.T. analyzed Visium ST data. H.H., A.Z an L.Z. interpreted all data. H.H. assembled figures and wrote the paper, with help from A.Z. and input from all authors.

## Methods

### Mice and ethics

Adult wild-type C57BL/6JOlaHsd mice (6-8 weeks, females) were purchased from Envigo and were group housed at the Technion SPF Pre-Clinical Research Authority under standard conditions in a 12-h light:dark cycle, at ambient temperature 21-23°C and and 30-40% relative humidity. Standard chow and water were provided *ad libitum*. All animal procedures followed the legislation under the National Institute of Health guidelines and were approved by the Institutional Animal Care and Use Committee of the Technion Israel Institute of Technology.

### Plasmid library generation ΔG-SAD-B19

A DNA fragment of 263bp was amplified with Pfu from vector D150pSADdeltaG-GFP-F2-ins1 using the primers: N21Bsi2: ccctcgctgctcgtacggtgnnnnnnnnnnnnnnnnnnnnnGTGTCAGATTATATCCCGCA N30Pst: GGCGATGGATCCTGCAGGTCC The amplified DNA from 8x 50µl PCR tubes was purified with 8x DFH300 silica column (Geneaid Biotech). The total yield was 3.4µg. 486ng of the purified PCR fragments where digested with BsiWI and PstI and cloned into the corresponding restriction sites of 18µg plasmid D151pSADdeltaG-GFP-F2-ins2 in a total volume of 60µl. Chemical competent E. coli cells (XL1-Blue MRF’ Kan, Agilent) were prepared by the method of Inoue *et al*. (Inoue et al. 1990). To 60 tubes with 50µl cells 1µl ligation reaction was added, incubated for 5min, heat shocked at 42°C and regenerated for 20min with 450µl SOC medium. An aliquot of the collected transformation reactions was used to calculate the complexity of the library, which was around 812.000 transformated cells. The cells from one plate were harvested with 5ml TB-medium and grown in 100ml TB-medium for 6h to an OD600 of 1. About 7.5mg DNA was isolated from 2L cells with the Megaprep Kit (Macherey Nagel).

### Barcoded ΔG-SAD-B19 rabies virus generation

Pseudotyped Rabies virus particles were prepared at the Viral Core Facility of the Charité – Universitätsmedizin Berlin (vcf.charite.de) using standard protocols (Osakada and Callaway 2013). Briefly, in order to recover G-deleted rabies virus from cDNA, B7GG cells (gift from Edward Callaway, Salk Institute) were transfected with a mix of the rabies virus genomic vector D151pSADdeltaG-GFP-F2-ins2 (Barcode vector BR-16) or D151pSADdeltaG-BFP-F2-ins2 (BR-17) (internal numbering), pcDNA-SADB19N (Addgene plasmid # 32630), pcDNA-SADB19P (Addgene plasmid # 32631), pcDNA-SADB19L (Addgene plasmid # 32632) and pcDNA-SADB19G (Addgene plasmid # 32633). All vectors were gifts from Edward Callaway. Transfection was performed with Lipofectamin 3000 (ThermoFischer Scientific).

### BR-16.1

Three days after transfection of three 75cm^2^ flasks (80% confluency), cells were transferred into three 175cm^2^ flasks. After 3 to 4 days supernatant was collected and 15ml per flask were used for infection of another three 175cm^2^ flasks (80% confluency). This was repeated 4 more times to amplify the virus. Finally, five 175cm^2^ flasks (80% confluency) were used for the infection with the 7ml amplified virus supernatant. After inoculation of further 4 days the supernatant was harvested and filtered through a 0.45µm filter.

### BR-16.2 / BR-16.3

Three days after transfection of three 75cm^2^ flasks (80% confluency), cells were transferred into three 175cm^2^ flasks. After 3 to 4 days supernatant was collected and 20ml per flask were used for infection of another two 175cm^2^ flasks (80% confluency). This was repeated 2 more times with four 175cm^2^ flasks to amplify the virus. After inoculation of further 4 days the supernatant was harvested and filtered through a 0.45µm filter.

### BR-16.4 / BR-16.5 / BR-16.6 ***and*** BR-17.1

Three days after transfection of ten 75cm^2^ flasks (80% confluency), cells were transferred into ten 175cm^2^ flasks. After 4 days supernatant was collected filtered through a 0.45µm filter and concentrated by ultracentrifugation (22K rpm, SW32 rotor, Beckmann). 2ml per flask were used for infection of another four 175cm^2^ flasks. After inoculation of further 4 days the supernatant was harvested and filtered through a 0.45µm filter. Production of BR-17.1 was identical, except that GFP in the rabies virus genome vector used had been replaced with BFP.

For pseudotyping the rabies with the envelope protein EnvA of the Avian Sarcoma and Leukosis virus (ASLV) the obtained supernatant containing unpseudotyped viruses was applied onto ten 175cm^2^ flasks containing BHK-EnvA cells (90% confluency) (gift from Edward Callaway, Salk Institute) for 5h followed by a transfer of the cells into new flasks with fresh media. After one day, media was changed a second time. After further 2 days EnvA-pseudotyped rabies viruses containing supernatant was collected, filtered and concentrated by ultracentrifugation. Rabies virus titer was determined by infecting a serial dilution of the virus in HEK293T-TVA cells (gift from Edward Callaway, Salk Institute). The efficiency of pseudotyping was tested *in vivo,* where we confirmed no infection by BC-ΔG-RV without prior helper virus administration (Fig. S1).

### Quality control barcoded ΔG-SAD-B19 *in vitro*

Barcodes counting is based on bulk amplicon sequencing. Briefly, HEK-TVA cells were infected with barcoded ΔG SAD-B19. 48hr later, ∼20% of the cells were infected with rabies. Cells were then trypsinized and aliquoted into 10^5 cells per vial. Total RNA was extracted from 5 vials of cells with the RNA extraction kit (NucleoSpin, Macherey Nagel). To count the barcode number of each library, we reverse transcribed the RV-*M*-gene, using an *M*-complementary primer with the Trueseq Read 1 sequence, and a 12bp UMI (T1-UMI-RV_*M*). First-strand was carried out on 40ng total RNA, in a 20μl reaction (20U/μL Maxima H Reverse Transcriptase (Thermo), 2.5mM dNTP, 1μM T1-UMI-RV_M, 1M Betaine, 6mM MgCl2 and 2U/μL RNAse Inhibitor), incubated 45min at 50°C, 5min at 85°C, then 4°C; followed by clean-up with 2x SPRI beads, and eluted in 10μl EB. Next, the RV barcoded region was amplified with primers complementary to RV-*M* gene RV_M-T2-PCR-forward (introducing Truseq 2) and Truseq 1 (T1-PCR-reverse) in a 20μl reaction (1x KAPA, 0.5M Betaine, 0.5μM primer mix), 3min 98°C, then 16 cycles of 15sec 98°C, 20sec 60°C, 1min 72°C, finally 1min 72°C, followed by 1.2x SPRI beads cleanup, and elution in 10μl EB. Finally, sequencing adaptors P5 and P7 were introduced following the 10xGenomics dual-index setup: 20-50ng PCR product was added to library PCR mix (50% KAPA, 10% Betaine, 20% primer Dual index (Illumina)), total 20μl; 45sec 98°C, then 16 cycles of 20sec 98°C, 30sec 54°C, 20sec 72°C finally 1min 72°C. PCR was followed by double-sided SPRI clean-up (0.6x SPRI and 0.8x SPRI). Amplicon libraries were sequenced on Illumina NextSeq2000.

### Viral injections

In vivo barcoded monosynaptic tracing is based on two stereotactic injections for AAV helper virus and RV respectively as described previously. Briefly, surgeries were performed on naïve C57BL/6 mice, aged 6-8 weeks. Mice were anesthetized using isoflurane, and maintained on 1.8% isoflurane (SomnoSuite Kent Scientific). Anaesthetized mice were fixed onto the stereotactic rig (Neurostar). Optic ointment (Duratears) was applied on both eyes and body temperature was maintained at 37°C with a heating pad. We observed heart rate and oxygen saturation throughout the procedure. Craniotomy was performed after applying Lidocain/Bupivacaine to the skull for local pain relief, with a 200μm diameter drill. Virus was injected with pulled glass pipette at steady rate (25-50nl/min). After injection, we waited 8-10 min before withdrawing, to prevent reflux. Target coordinates were obtained from The Mouse Brain in Stereotaxic Coordinates, 3rd edition, Keith B.J. Franklin, George Paxinos. Buprenorphine (0.05-0.1mg/kg) was given for analgesia on the day of surgery and the day after. 21±4 days after AAV injection, RV was injected to the same target, following the same surgical procedure. Brains were harvested 7±1 days after RV injection.

For the scRNA-seq and Visium ST experiments we used helper virus with the B19 optimized glycoprotein; ssAAV-8/2-hCMV-hHBbI/E-TVA_mCherry_2A_oG-hGHp(A). This virus was generated by Viral Vector Facility VVF Zurich (v534 serotype 8, VVF Zürich), based on pAAV-CMV-TVAmCherry-2A-oG; a gift from Marco Tripodi (Addgene #104330) (Ciabatti et al. 2017).

For all other surgeries, we used helper virus with the N2c-version of glycoprotein; ssAAV-2/1-Syn-H2B-mCherry-TVA-N2c.G-WPRE3. This virus was generated by the Viral Core Facility of the Charité Universitätsmedizin Berlin (Cat# BA-413, serotype 1), based on Addgene plasmids pAAV-EF1a-DIO-HTB (# 44187, gift from Edward Callaway) and pCAG-N2cG (# 100811, gift from Ian Wickersham).

Details for BC-ΔG-RV libraries are provided in the section “Barcoded ΔG-SAD-B19 rabies virus generation”.

### scRNA-seq of barcoded ΔG-SAD-B19 injected brains

To isolate single BC-ΔG-RV (GFP-library BR-16.1) infected cells for scRNA-seq, we modified a protocol for gentle adult neuron dissociation (Hochgerner et al. 2023). 8 days after bilateral stereotaxic injections to the CA1, one mouse was sacrificed by an overdose of Ketamine/Xylazine, and transcardially perfused with ice-cold carboxygenated NMDG-aCSF (Ting et al. 2014). The brain was extracted, then mounted and sectioned on a Leica VT1200S automated vibrating microtome, to 500μm slices. Fluorescent regions were quickly microdissected in ice-cold aCSF under a fluorescent stereomicroscope (Nikon SMZ18). Next, collected tissue pieces were digested in 800-100μl papain digest solution with DNase (Worthington Papain system, concentrations as recommended by manufacturer, but vials were reconstituted in aCSF), 25min at 35-37°C. Starting 15min into incubation, tissue was gently aspirated in wide-bore fire-polished Pasteur pipettes and returned to incubate, until most tissue easily broke apart. We filtered the suspension through a 30μm strainer (CellTrix, Partec) to ice-cold microtubes coated with 30% BSA, and pelleted cells at 4°C, 5min, 200g. Supernatant was fully removed, cells immediately resuspended in 500μl fixative (DSP and SPDP in sodium phosphate buffer pH8.4) (Gerlach et al. 2019), and incubated 40min, at RT, gently shaking. The fixation was stopped with 1 volume quenching buffer (0.1M Tris-HCl 150mm NaCl, pH7.4) 10min, fixed cells pelleted, and resuspended in PBS with 0.3%BSA and 1% recombinant RNase inhibitor (RRI, Takara Bio). Next, we enriched for GFP+ cells on a FACS Aria II (BD), or Bigfoot Spectral Cell Sorter (Thermo Fisher). ∼19k sorted cells were immediately processed for encapsulation and scRNA-seq on the 10xChromium NextGEM, in two samples, following the manufacturer’s instructions. Libraries were sequenced on the Illumina NovaSeq NGS platform, at a depth of ∼60K reads per cell.

### Histology and ROInet-seq

#### Histology

7d after RV injection, mice were sacrificed after an overdose of Ketamine/Xylazine, and transcardially perfused with PBS followed by 4% paraformaldehyde (PFA). Brains were dissected, postfixed in 4% PFA overnight and cryoprotected in 30% sucrose solution with PBS for 24 hr. After OCT embedding, brains were cryosectioned full-volume to 50μm floating sections, mounted to Superfrost Plus slides (Epredia), counterstained (NucBlue, Thermo) and coverslipped (Fluoromount-G, Thermo). Images were taken using an epifluorescence microscope (Nikon Eclipse Ti2) at 4x magnification. Slides were stored at -80 °C until further use.

#### ROI microdissection andlysis

For ROInet-seq, coverslips were gently removed in PBS, and under a fluorescent stereoscope, ROIs (tissue with RV-tagged cells) were microdissected on ice into stripes filled with lysis buffer (30mM Tris-HCL pH 7.5, 10mM EDTA, 1% SDS, 0.1μg/μl Proteinase K). Dissected tissue was vortexed vigorously and lysed for 1h at 50°C, followed by 10min at 70°C, then placed on ice. Lysates were stored at -80°C until further use.

#### Bead preparation and polyA capture

To capture all polyA-transcripts, introduce a 12bp UMI, 16bp ROI-specific indexes, and Truseq read 1, we loaded RT-primer (T1-ROI_index UMI-OdT) to Streptavidin beads, as follows. First, Streptavidin-C beads (5μl per ROI, Thermo) were washed twice with 2xBWT buffer (10mM Tris-HCl pH 7.5, 1mM EDTA, 2M NaCl) and resuspend with equal volume of 2xBWT buffer. In separate wells, 3μl of each unique RT primer (T1-ROI_index UMI-OdT) were added to 5μl washed beads, completed with UPW to total volume of 10μl, mixed and incubated for 20min in room temperature, 1200rpm. After incubation, RT primer-loaded beads were magnetically immobilized and washed with TNT (20mM Tris-HCl pH 7.5, 50mM NaCl, 0.02% Tween 20) and resuspend with Capture mix (50μl per ROI_index: 2xBWT, 0.5U/μl RRI (Takara)). Equal volume of lysate was added to capture mix and incubated for 1min at 56°C followed by 20min incubation at room temp, shaking 2000rpm. After incubation beads were placed on a magnet, supernatant was removed and beads were washed thoroughly 3x with ice-cold LiWT buffer (10mM Tris-HCl pH 7.5, 1mM EDTA, 150mM LiCl) and resuspend with 10μl UPW (suitable for 3 reactions). RT primers with captured polyA-transcripts were released from beads by 2min incubation at 80°C, and transferred. Samples were stored at -80°C until further use.

#### Reverse transcription

Reverse transcription was carried out on 2.7μl sample, by adding RT mix (1x Maxima buffer, 20U/μl Maxima enzyme, 1M Betaine, 2mM dNTPs, 2U/μl RRI, 20mM DTT, 6mM MgCl2) to a total reaction volume of 6μl, and incubating 45min at 50°C, 5min at 85°C, then 4°C.

#### Clean-up

After RT, samples were cleaned up using 2x volumes CleanUp buffer (5 : 1 volumes of SPRI : Guanidine Thiocyanate 6M solution) by 10min incubation followed by 2x 30sec 80% ethanol wash using bead magnet, 1min air dry and elution with 7.4μl elution solution 1 (1x Reducing Agent B (10xGenomics), 0.01% Tween-20 in buffer EB (Qiagen)).

#### PCR amplification

The RV barcoded region was amplified with primers complementary to RV-*M* gene (introducing Truseq read 2) and Truseq read 1, as follows. The eluate was added to 12.6μl PCR mix (1x KAPA HS, 0.5M Betaine. Total reaction volume 20μl) with 1μM primers (RV_M-T2-forward and T1-PCR-reverse), for targeted amplification of the RV M-gene with BC; 3min 98°C, then 17 cycles of 15sec 98°C, 20sec 60°C, 1min 72°C, finally 1min 72°C. After PCR, samples were cleaned with 1.2x SPRI protocol and finally eluted in 10μl EB. Sample concentration was evaluated using Qubit.

#### Library indexing and sequencing

Finally, sequencing adaptors P5 and P7 were introduced following the 10xGenomics dual-index setup: 20-50ng PCR product was added to library PCR mix (50% KAPA, 10% Betaine, 20% primer Dual index (Illumina)) total 20μl; 45sec 98°C, then 16 cycles of 20sec 98°C, 30sec 54°C, 20sec 72°C finally 1min 72°C. At this point, all ROI samples were pooled to one tube, followed by double-sided SPRI clean-up (0.6x SPRI and 0.8x SPRI). Concentration was evaluated using Qubit and fragment size on Bioanalyzer or TapeStation. The final ROIs amplicon libraries resemble 10xChromium v3/NextGEM sequencing libraries and were sequenced at a depth of ∼10M reads per sample (one brain) on Illumina NextSeq2000.

In its final implementation, a typical ROInet-seq experiment was microdissected and processed for RNA-sequencing within 2 experimental days (day 1, microdissection and lysis; day 2, molecular biology processing); when preceded by ∼1 day for careful brain alignment and ROI-dissection strategy design. PCR amplification introducing the final library indexes can in principle be carried out with all ROI samples in pool, as demonstrated for the experiment presented in Fig. S4. This saved ∼2h of manual handling time but increased representation of noisy BCs; requiring more stringent computational processing, detailed in “ROInet-seq data processing and analysis”.

### ROInet-seq data processing and analysis

ROInet-seq fastq files contained ROI-index, UMI (both Read 1) and the target sequence on the 3’UTR of the RV-*M*-gene, including the BC-read (Read 2). First, we aligned ROI-index reads with the known list of ROIs wells that were collected. In Read 2, we called the BC read by aligning the known, permanent sequences flanking the BC in the *M*-3’UTR, and extracting every variable 21nt sequence inside. Reads per ROI were restricted to 1000 per fluorescent cell detected in that ROI. This was necessary since sequencing depth per ROI could not be accurately controlled in the pooled setup. Only BCs that were detected in the *in vitro* library (>10 reads in HEK-TVA culture, see “Quality control barcoded ΔG-SAD-B19 in vitro”) were included in analysis. We filtered BCs with low confidence, by removing BCs of below 50 reads over all ROIs. Any BC detected below a threshold of 10UMIs per ROI was also removed.

Next, we restricted network analysis to the most widely spread unique BCs as follows: Unique BCs were ranked by network size (i.e., number of ROIs where the BC was detected; Fig. SX)). We selected for the BCs that were higher in either: (a) the detected ‘knee’ of this ranked BC list +30%, or (b) the number of counted starter cells plus 50%. This was done to remove potentially dubious BCs, based on the observation that individual BC-ΔG-RV-infected cells in scRNA-seq (Fig. S2) and Stereo-seq ST (Fig. 6, Fig. S11) contained no more than ∼1.5BCs/ starter cell. BC detected above these thresholds were included towards the final BC count matrix (Table S1), used to plot overlaps of BCs between ROIs, and global or single-barcode networks.

Finally, to avoid the risk of multicell-infection of identical BCs, we removed BCs that were highly abundant in the *in vitro* library (>10^-4^ in HEK-TVA) or in all three *in vivo* experiments (>10^-4^ in 98-1, -5 ***and*** -6).

### Stereo-seq cell-resolved spatial transcriptomics networks reconstruction

#### Tissue collection and spatial transcriptomics

Transcardial perfusion with aCSF was performed under deep anesthesia on the harvest day. Brains were snap frozen on dry ice and stored at -80°C until sectioning the following day. Tissue mounting was performed according to the steps outlined by the manufactorer (STOmics, Stereo-seq transcriptomics set for chip-on-a-slide V1.2, manual version A1). The tissue was sectioned into 10μm-thick slices, and the target region (hippocampus) was trimmed with a blade and transferred onto Stereo-seq transcriptomics chips, such that each chip contained nine consecutive hippocampal sections. The anterior-posterior distance between every two consecutive sections was 50μm. Subsequent steps, including image acquisition, tissue permeabilization, reverse transcription, cDNA release and amplification, and library preparation, were all carried out according to the protocol.

#### Data analysis

To calculate the expression of rabies per spot, we summed the number of detected molecules for rabies transcripts *GFP*, *M* and *P*. Spots with detected RV-expression that had at least 5 neighbors in a radius of 20 spots (∼10μm) were retained for segmentation of RV cell somas. We next performed spatial clustering using DBSCAN (epsilon 15, minpoint 5). This resulted in 1893 clusters that were putative RV-infected cells and were segmented. We further subdivided clusters larger than 4000 pixels using k-means (*k* = #of pixels / 4000; rounded to nearest greater integer). Then, clusters smaller than 100 were discarded. Next, all BC reads with x-y position within the bounds of remaining clusters were assigned to these clusters. When the highest represented BC within a cluster was not detected by at least 3 reads, this cluster was discarded, too. The remaining clusters were considered as putative network cells (n=774); by their RV transcript expression, cell size and BC detection, and are seen as yellow outlines (Fig. 6) or black outlines (Fig. S11). All clusters that were discarded according to these criteria are seen as blue outlines in Fig. 6 (n=1119). Finally, all BCs supported by just 1 read per putative network cell were discarded. All remaining BCs are analyzed as networks and presented in Figs. 6 and S11.

### Visium ST networks sequencing and analysis

#### Tissue collection and spatial transcriptomics

Transcardial perfusion with aCSF was performed under deep anesthesia on the harvest day. Brains were snap frozen on dry ice and stored at -80°C until sectioning the following day. The brain was cryo-sectioned into 20μm, by cutting only the hippocampus region and collecting to capture areas on the Visium Spatial Transcriptome slide (Fig. S6). The gap between two consecutive sections collected to the capture area was 100μm. All adjacent sections were collected to a SuperFrost Plus Slide (Epredia) and stored at -80°C until processing. Visium ST slides were processed as per the manufacturer’s instructions, with the following specifications: Slides were fixed with methanol, then DAPI counterstained and imaged on an inverted epifluorescence microscope (Nikon Eclipse Ti2). Tissue was permeabilized for 30 min. The mRNAs from the sections were then captured by the primers including UMI, spatial barcode and sequencing primer on the Visium ST slides. After the RT reaction and second strand synthesis on slides, the cDNA library was transferred into tubes, fragmented and indexed. For detection of RV-barcodes and AAV transcripts, cDNA was additionally amplified in two separate PCR reactions, with RV_M-T2-PCR-forward / T1-PCR-reverse (RV-*M*-gene) or AAV-T2-PCR forward / T1-PCR-reverse (AAV-*oG*-gene), using 20-30ng cDNA, each in a 10μl reaction (1x KAPA, 0.5M Betaine, 1μM primer mix), 3min 98°C, then 8 cycles (RV-*M*) or 10 (AAV-*oG*) cycles of 15sec 98°C, 20sec 53°C, 1min 72°C, finally 1min 72°C, followed by 1x SPRI beads cleanup, and elution in 4μl EB. Finally, sequencing adaptors P5 and P7 were introduced following the 10xGenomics dual-index setup: Eluted PCR product was added to library PCR mix (50% KAPA, 10% Betaine, 20% primer Dual index (Illumina)), total 20μl; 45sec 98°C, then 12 cycles of 20sec 98°C, 30sec 54°C, 20sec 72°C finally 1min 72°C. PCR was followed by double-sided SPRI clean-up (0.6x SPRI and 0.8x SPRI). Amplicon libraries were sequenced to 140-260M reads per sample on Illumina NextSeq2000.

#### Data analysis

The methodology for analyzing Visium data and detecting barcodes is implemented through a comprehensive MATLAB script designed for the integration of rabies barcode information with spatial context derived from Visium experiments. It proceeds to process barcode data from distinct experimental conditions, including Vis11_A1, Vis11_B1, Vis11_C1, and Vis11_D1, as well as RFP-AAV SAM files. This involved organizing data into: cell IDs, unique sequences, and read frequencies, leading to the creation of a unified table combining rabies and AAV data. Further, since sequencing depth of individual capture spots was unequal, we next normalized spots to each other; accounting for the total number of reads per spot, and the number of BCs detected in that spot, as follows: For each spot, the total read count was divided by the square root of the number of unique BCs in that spot. Finally, we filtered noisy BCs, by removing BCs that were below an average expression threshold or had null empty entries. Genes and BCs shown in the Figures were plotted in a binary fashion, i.e., each spot with a detected gene or BC is shown positive. We further incorporate functionality to subset and process specific barcode sequences from FASTQ files, saving the resulting subset and identifying common elements with loaded barcodes. Additionally, we used DAPI images of the Visium ST sections to map each section in its anatomical context by semi-manual aligning sections to the Allen Reference Atlas—Mouse Brain (atlas.brain-map.org) and correcting for distortion caused by sectioning along the dorsoventral and mediolateral axes. Sections were on average 100μm apart. mRNA-capture spot diameters were 55μm, centers 100μm apart (x–y resolution). 18,177 capture spots covered the tissue and contained on average ∼3,300 genes and ∼9,900 UMIs. 1848 of the capture spots were positive for RV (expressing positive values higher than one). We called the BC read by aligning the known, permanent sequences flanking the BC in the *M*-3’UTR and extracting every variable 21nt sequence inside. We finally corrected for sequencing errors by pooling any BCs with Hamming distance of ≤3 with its most abundant version.

### osmFISH

To detect viral transcripts in sections adjacent to the ones sampled for Visium ST, we performed in situ hybridization according to osmFISH (Codeluppi et al. 2018), with minor modifications. We collected 10μm sections to Superfrost Plus slides (Epredia) and stored them at -80°C until processing. After the sections were balanced at RT, the tissue was fixed in 4% PFA for 10 min, washed 5 min with 2x SSC buffer, and

cleared with 8% SDS for 30 min. Tissue is incubated with hybridization mix (2x SSC, 10% dextran sulfate, 10% formamide, 1mg/ml E. coli tRNA, 2mM ribonucleoside vanadyl complexes, 0.2ml/ml bovine serum albumin) without probes for 5 min at RT and then 4h at 38.5 °C in hybridization mix with probes (250nM). After wash with 20% formamide in 2x SSC, slides were mounted (ProLong Diamond Antifade, Thermo) and imaged at 10x magnification, on an inverted epifluorescence microscope (Nikon Eclipse Ti2).

### Oligonucleotide DNA sequences

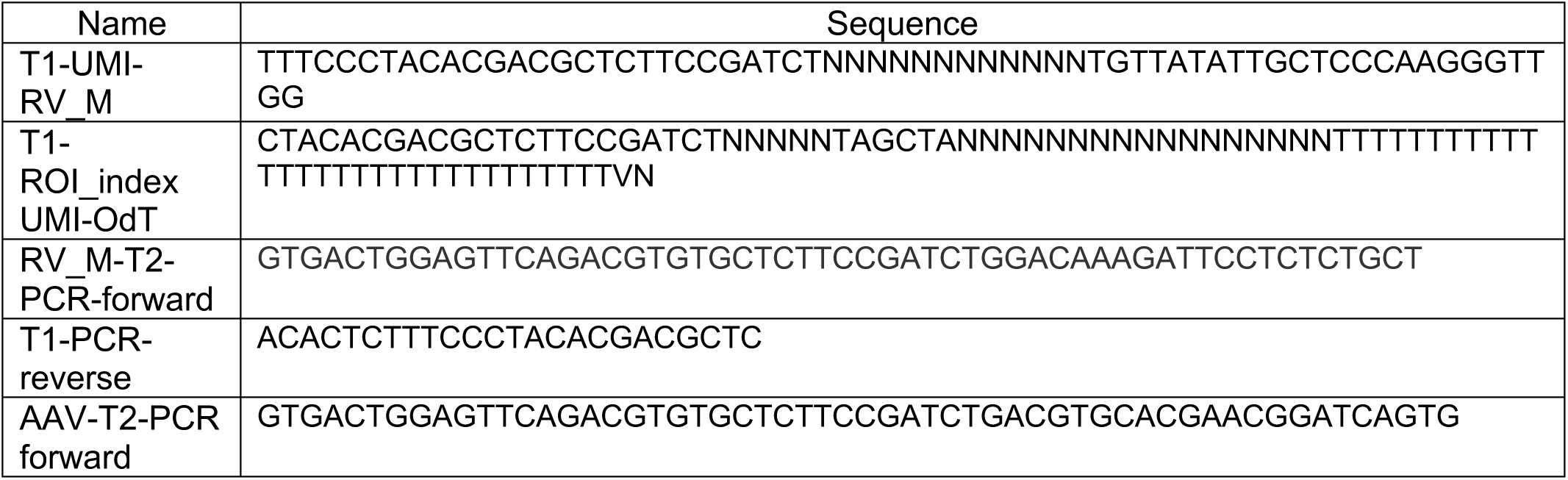

### Data availability

The raw sequencing data generated in the current study will be made available through NCBI GEO. Source Data (BC count tables for ROInet-seq experiments) are provided as supplementary files with this paper. For convenience, we additionally provide single-cell expression dataset with annotations and metadata as a table at figshare, made available during review.

### Code availability

Custom code used to perform the analysis will be made available during review.

