## Supplementary material for "Brain-wide monosynaptic single-neuron connectivity with ROInet-seq": Supplemetary Information

**Table S1, Barcode count matrix SSp.** Related to Fig. 3. Barcode counts per ROI, for SSp ROInet-seq experiments 98-1, 98-5, 98-6. This table contains all BCs before filtering abundant BCs (Fig. 4, see Methods).

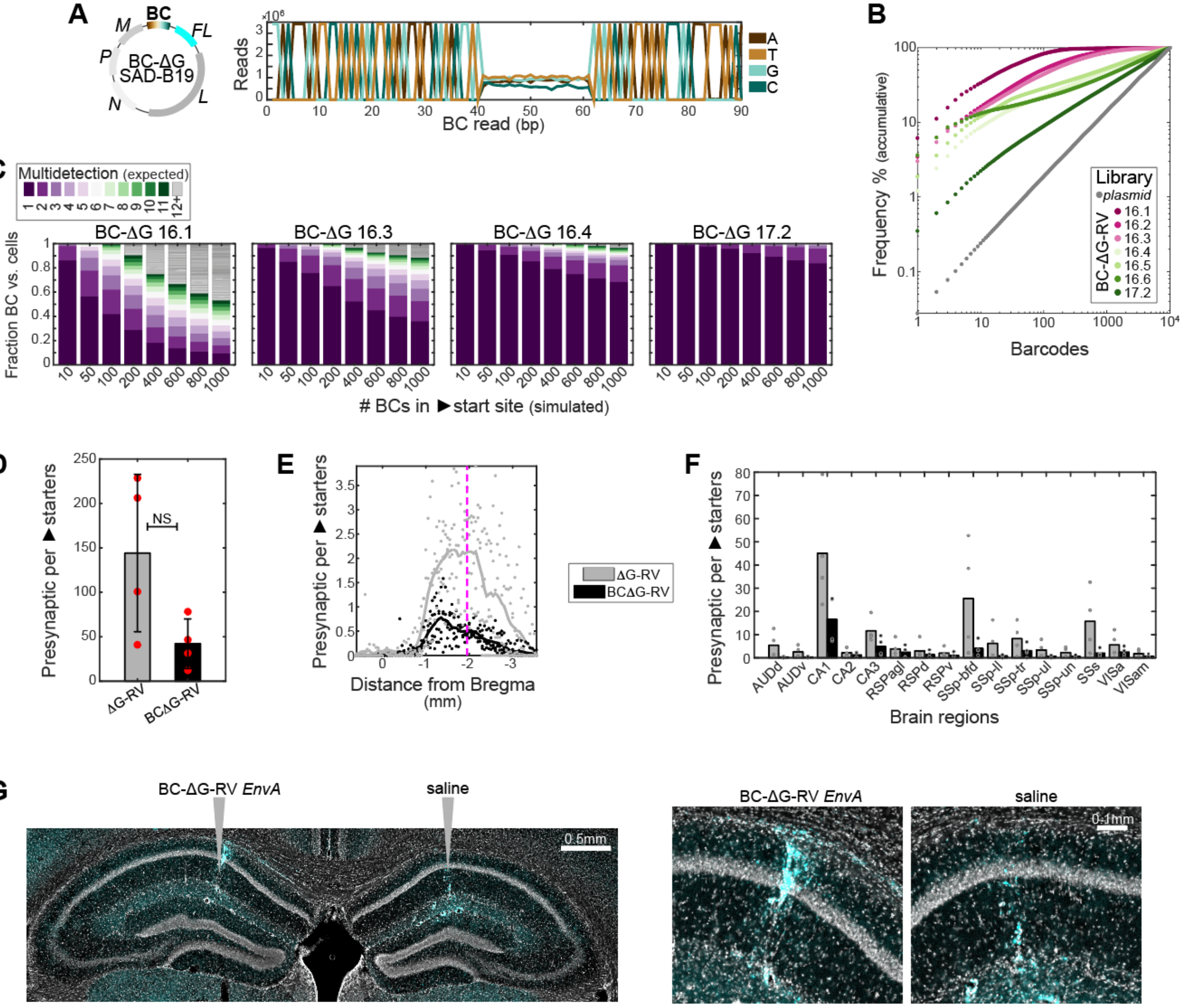

**Fig. S1, Characterization of BC-ΔG-RV libraries.**

(**A**) Sequencing verification of a uniformly random 21nt BC insert, shown as pseudo-Sanger bp frequency plot.

(**B**) Accumulative frequency of top 10k most frequent barcodes per RV-library, vs. the optimally uniform plasmid library.

(**C**) Expected detection of a unique BC in 1-12+ cells, given 8 scenarios based on 10-1000 starter cells, per library.

(**D**) Quantification of presynaptic-per-starter cells after infection to the CA1 (unilateral) with ΔG-RV (n=4; grey; 144.1 +/- 88.5) and BC-ΔG-RV (n=4; black; 42.2+/-27.9).

(**E**) As (**D**), quantification of presynaptic-per-starter neurons resolved to the sections along the anterior-posterior axis. Pink line, injection coordinates.

(**F**) As (**D**), quantification of presynaptic-per-starter neurons, summed by brain region.

(**G**) *In vivo* EnvA pseudotype test. Without prior infection with AAV helper virus, we unilaterally injected either BC-ΔG-RV (left) or saline (right). Both injections show signs of tissue damage along the injections' trajectories (increased autofluorescence), but no labelling of neuronal soma.

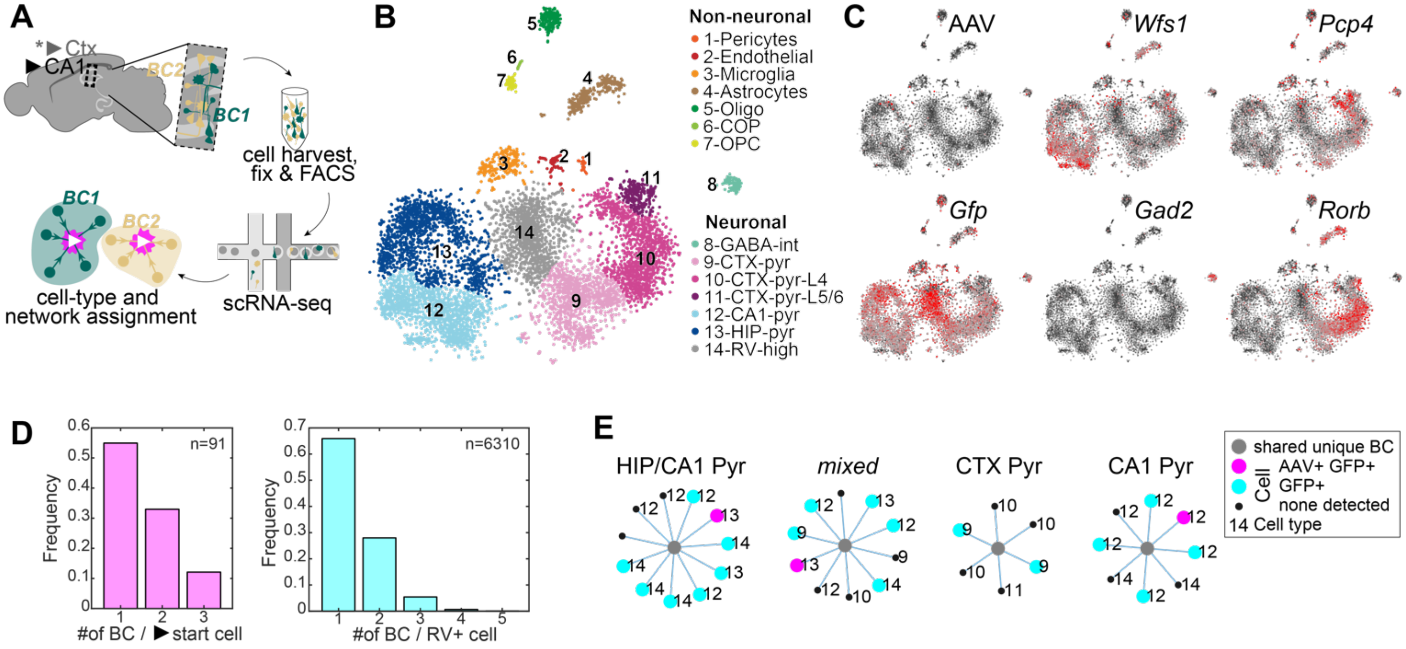

**Fig. S2, Single-cell RNA-seq on BC-ΔG-RV-infected neurons *in vivo*.**

(**A**) Experimental schematic, BC-ΔG-RV infection of hippocampus CA1, with secondary infection in isocortex dorsal to CA1, followed by dissociation, fixation, FACS and scRNA-seq; to simultaneously detect transcriptomes and BCs.

(**B**) tSNE visualization of 8032 cells, colored by cell type. Among 6979 neuronal transcriptomes, 129 were GABAergic interneurons, 2744 had a predominantly cortical pyramidal signature (1566 of which L4/5/6) and 2937 a hippocampal pyramidal signature (1418 of which CA1).

(**C**) Expression of 4 neuronal genes, *Gfp* (RV) or any one AAV-gene (*TVA, oG, mCherry*) in all cells visualized in (**B**). Grey, low; red, high; black circle, 0. Note, cluster 14 featured the highest expression of *Gfp* (RV), but overall low-quality transcriptomes with general neuronal markers.

(**D**) Frequency of detecting single or multiple BCs in a single cell; in starter cells (91 neurons, as in Fig. 1F) or all BC-ΔG-RV+ cells (total, 6310 BC+ neurons).

(**E**) Four examples of potential single-cell networks; i.e., cells sharing a unique barcode. Over all networks, most have no verified starter cell (cell with AAV and GFP detected), and most have some degree of mixing of transcriptomic types from CTX and HIP, as shown in one example.

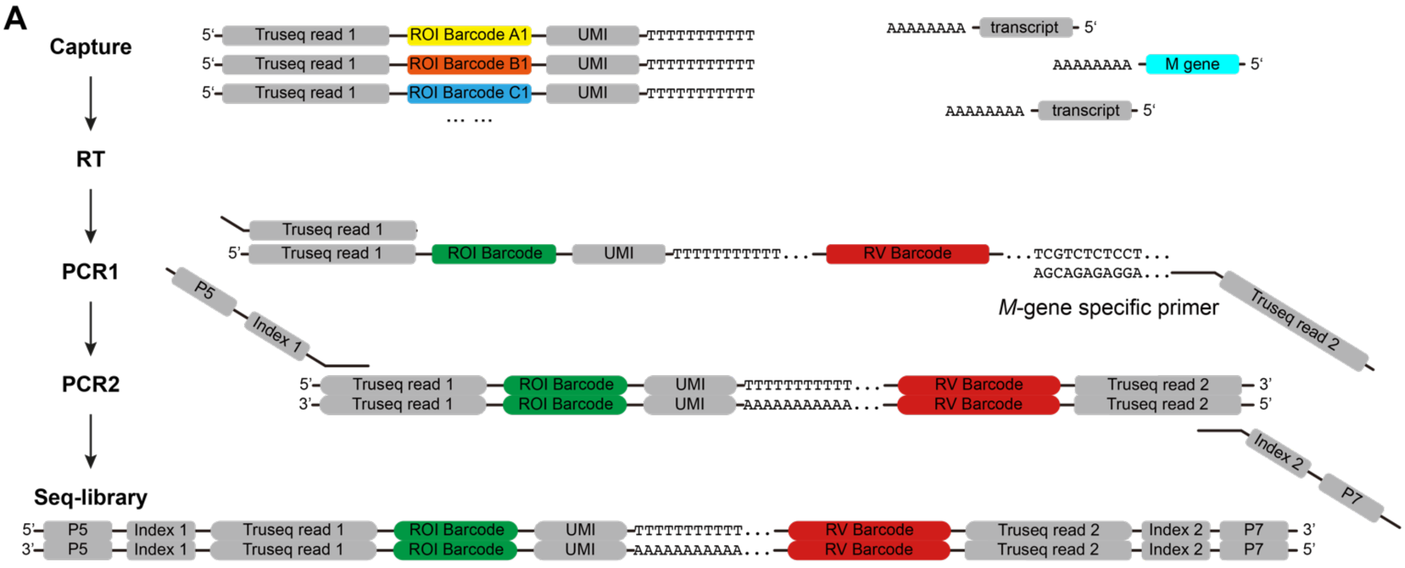

**Fig. S3, Molecular biology workflow for ROInet-seq.**
Schematic of RT and BC-specific amplification from fixed tissue ROIs in ROInet-seq. Each ROI is collected to a separate well, loaded with lysis buffer and polyT primer carrying a unique ROI barcode (spatial index). In reverse transcription (RT), the indexed polyT captures polyA transcripts present in that ROI, including RV-genes such as *M* (with the network BC in its 3'UTR). cDNA is amplified in two consecutive PCR reactions, first using an oligo specific to a sequence flanking the *M*-gene (and BC), and then sequencing library prep to introduce library indexes and P5/P7.

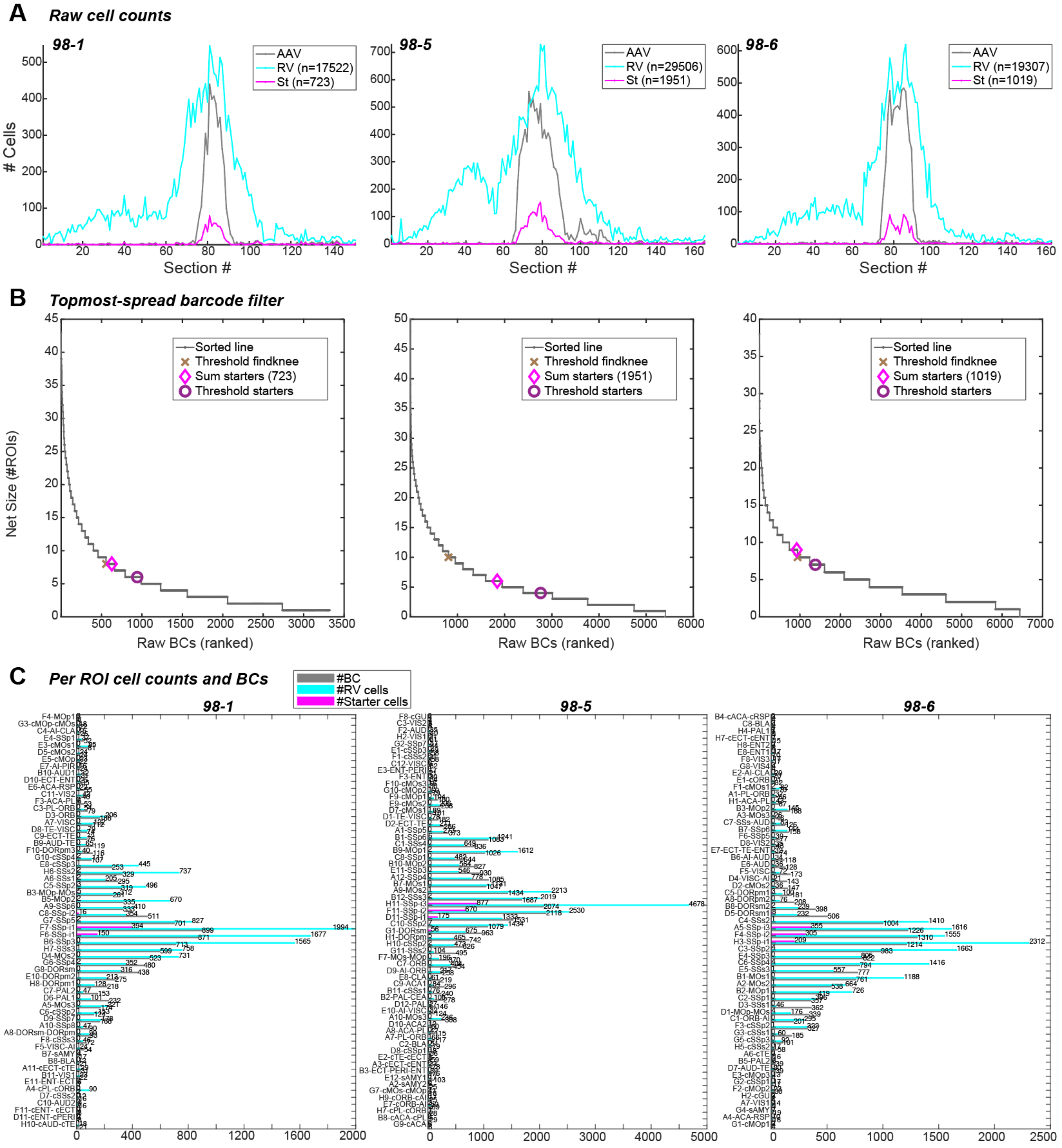

**Fig. S4, ROInet-seq data processing – cell counting and BC detection.**
(**A**) After BC-ΔG-RV infection of the SSp, automated cell detection of AAV+ and RV+ cells across the full brain in anterior-posterior section order (50μm, gapless collection). Fluorescent overlap of signals (AAV+RV+) is considered a starter cell (St).
(**B**) Ranking of BCs by their detection in number of ROIs, over all BCs. Marks along the sorted line indicate thresholds used to select the topmost widely spread BCs for network analysis, where higher threshold was applied (see Methods).
(**C**) Summary of all ROIs per experiment, with imaging-based cell counts and BC detection.

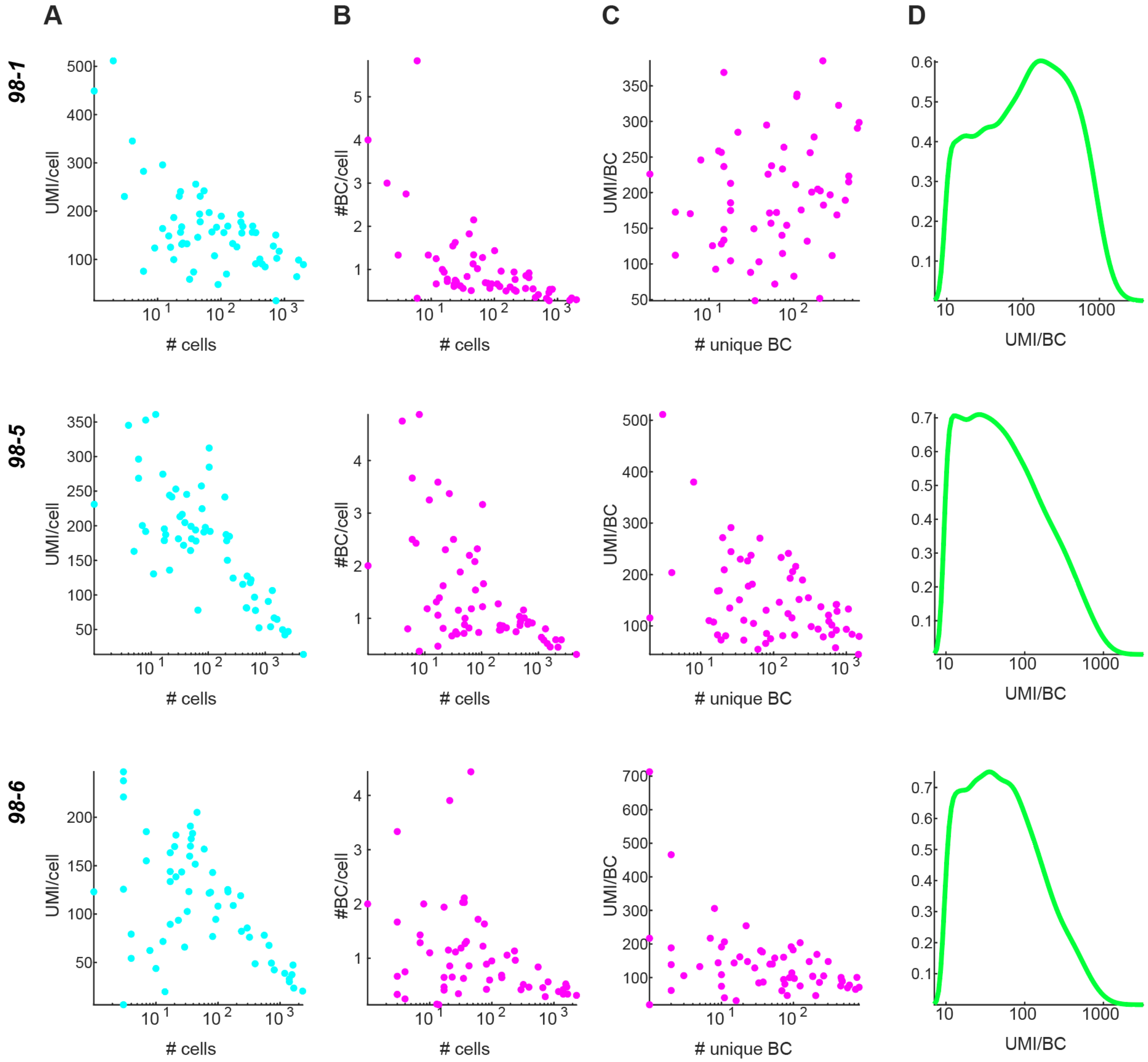

**Fig. S5, Per-ROI quantifications of cells, BCs and UMIs in ROInet-seq.**
Per experiment (row), where each ROI is one dot (**A**-**C**)
(**A**) Scatterplot of UMI/cell as a function of number of cells detected (imaging).
(**B**) Scatterplot of BC/cell as a function of number of cells detected (imaging).
(**C**) Scatterplot of UMI/BC as a function of BCs.
(**D**) Distribution of copies (UMI) detected per unique BC.

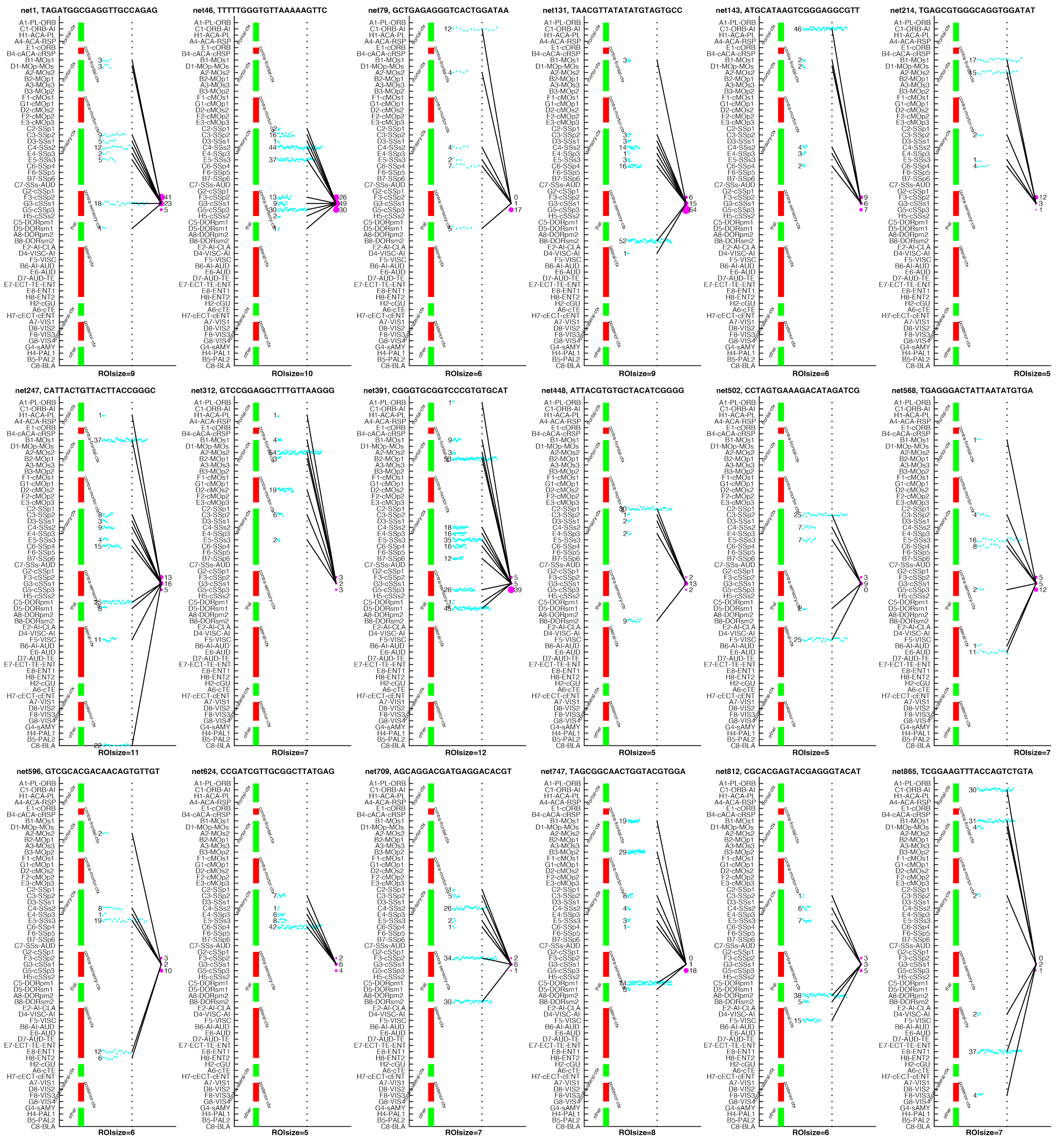

**Fig. S6, Input networks to single SSp neurons.**
Brain-wide input networks to 18 single SSp start site neurons in brain 98-6 at ROI resolution, visualized as dots, where each cyan dot represents detection of 10 unique network BC copies (UMI) in input ROIs, and magenta blob size represents unique BC copies in SSp start site.

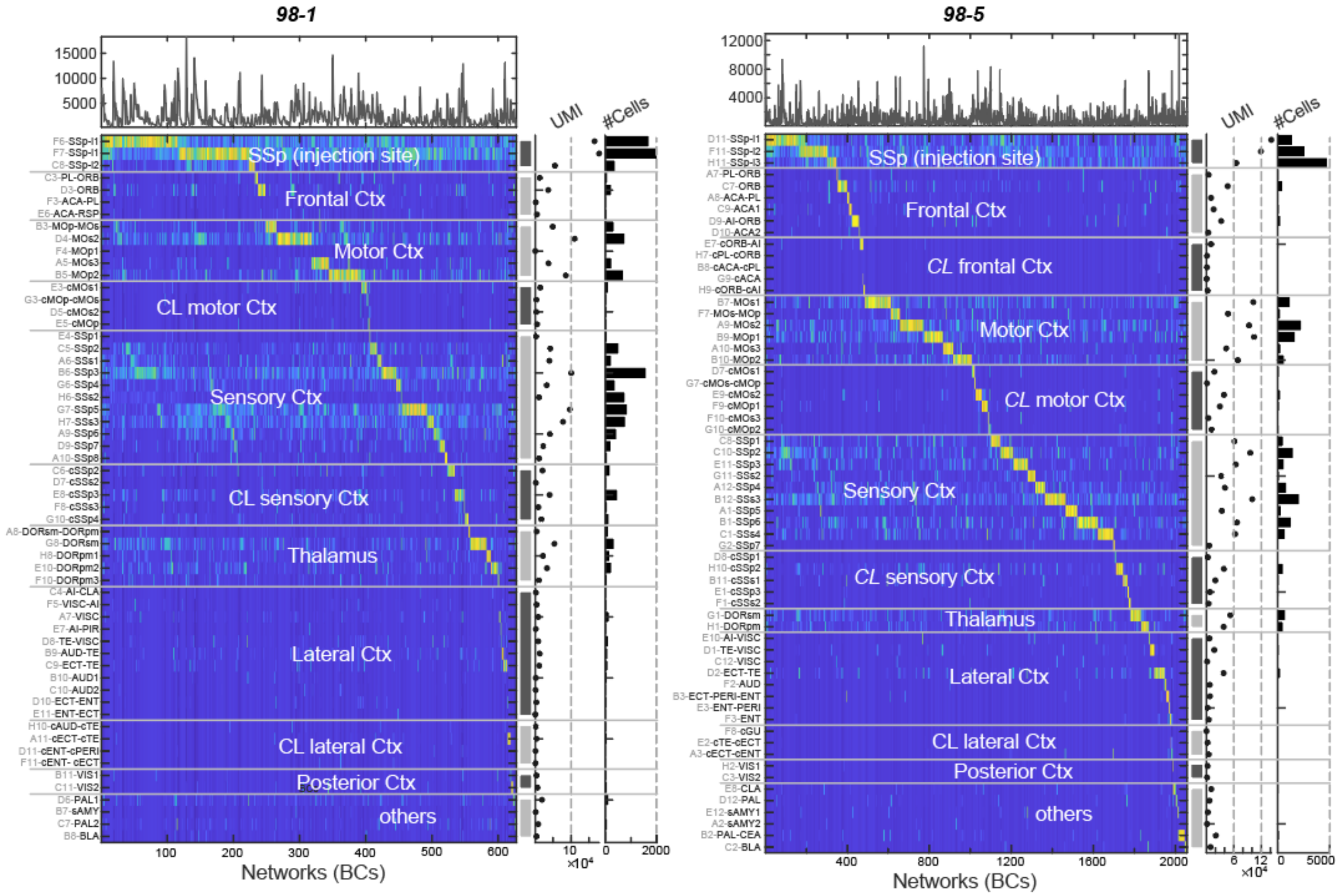

**Fig. S7, Brain-wide input networks of single cells in the SSp, detected by ROInet-seq.**
Single-cell networks across all ROIs in brains 98-5, 98-6, ordered by dominantly detected ROI. Colormap represents network BC copies (UMI) (blue, low; yellow, high).

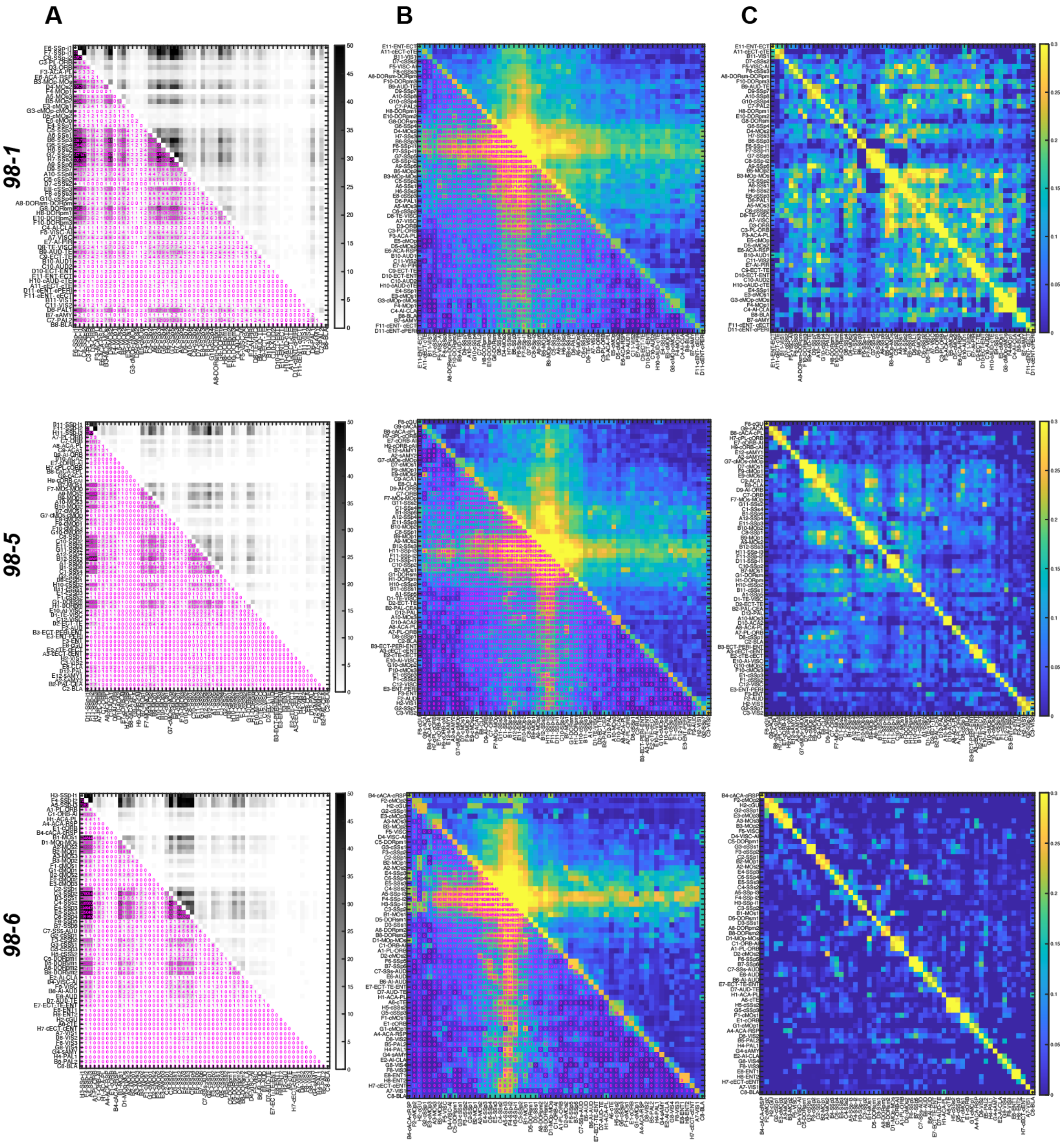

**Fig. S8 Brain-wide regional overlaps of SSp-cell network.**

Convergent regional inputs to single SSp start site neurons in brains 98-1, 98-5 and 98-6. Co-detection of network BCs between individual ROIs;

(**A**) Absolute overlap of all detected network BCs, in ROI order (as Fig. 5, S6);

(**B**) Absolute overlap of all detected network BCs, in order of similarity;

(**C**) As pairwise correlation, in order of similarity (as (**B**)).

Color-/greyscale represents min to max values normalized per ROI (excluding the diagonal).

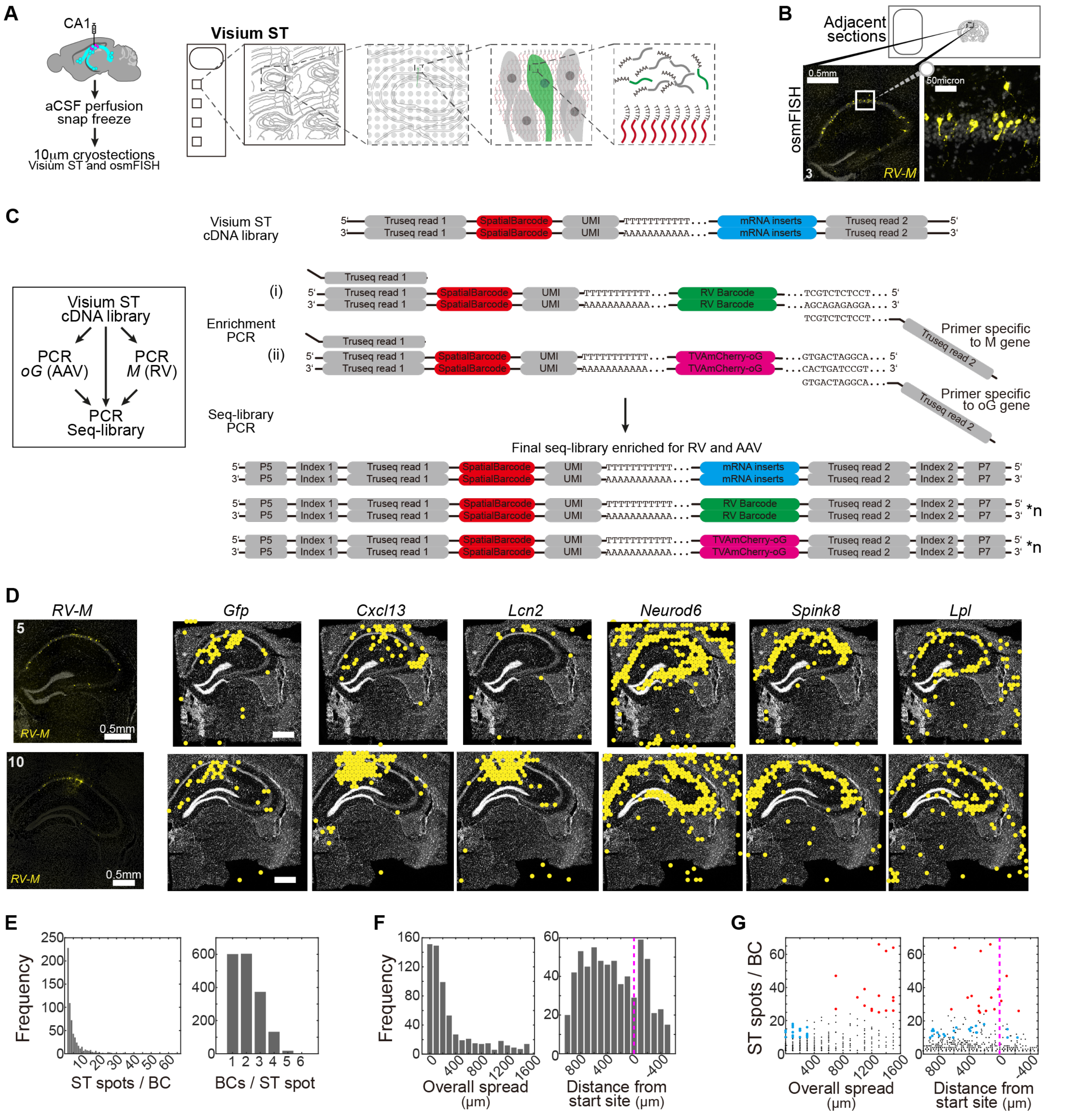

**Figure S9 Visium Spatial Transcriptomics (ST) for increased resolution of local network reconstruction.**
(**A**) Experimental outline Visium ST experiment: Collection of 16 coronal cryosections of the hippocampus, spanning the anterior-posterior axis at 100mm spacing. Per ST capture area, 4 sections were collected. After cDNA library construction, sequences of interests (AAV and RV) were enriched, and sequenced.
(**B**) Validation of RV-infection in sections consecutive to the Visium ST-sampling, using single molecule FISH (osmFISH) against the RV matrix gene (M). Example and zoom-in on osmFISH section adjacent to Visium ST section 3.
(**C**) Workflow for amplification of RV (BC) and AAV (TVA) -related transcripts, from full cDNA.

(**D**) Visium ST gene expression of RV (*Gfp*), pyramidal neuron (*Neurod6, Spink8, Lpl*) and inflammatory markers (*Lcn2, Cxcl13*) in sections 5 and 10; with single molecule FISH-validation of RV matrix gene (*M*) expression in consecutive sections shown on the left. Scale bars, 0.5mm.

(**E**) Detection of RV-barcodes (BCs) in Visium ST: left, detection of barcodes in ST spots, where most BCs were restricted to 5 spots; and right, number of BCs per spot, where most RV+ spots contained one or two barcodes.

(**F**) Left, distribution of total spreading distance of barcodes; and right, spreading distance relative to the start site.

(**G**) Left, scatter plot of quantified barcode detection, by total spreading distance, and right, scatter plot of quantified barcode detection, by relative position to the start site (average per BC). Every spot represents one barcode; red labels the top 16 most abundantly detected BCs, blue, 22 of the most locally restricted BCs.

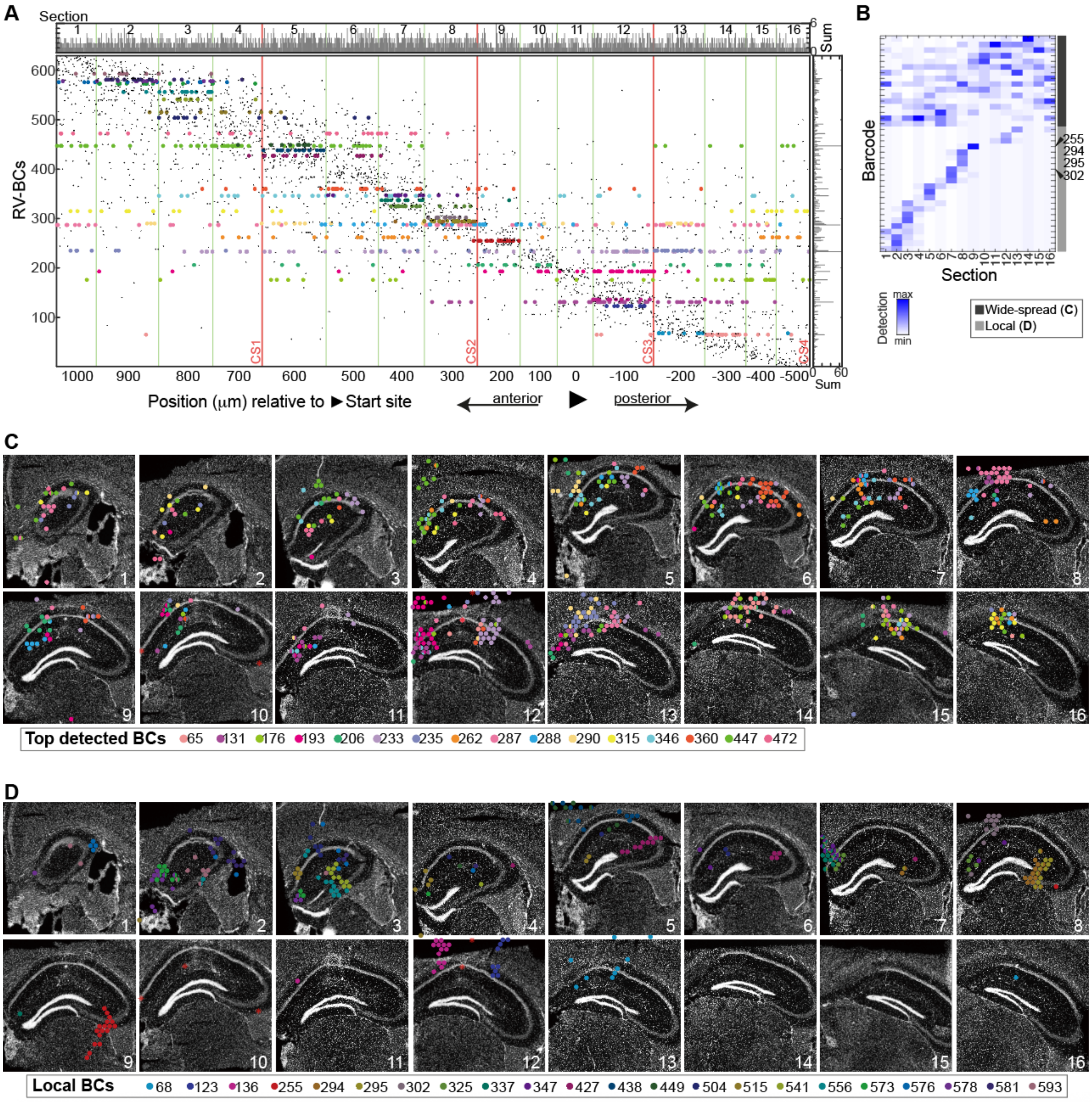

**Fig. S10 Visium Spatial Transcriptomics (ST) for local network reconstruction.**
(**A**) Distribution map of all 629 detected barcodes (rows), across all RV+ 1848 Visium ST spots (columns), across the 16 a.-p. sections.
(**B**) Detection of 16 top detected and 22 locally mapping BCs, across 16 a.-p. sections. Blue, high; white, low.
(**C**) Spatially resolved detection of the top 16 abundant barcodes, in 16 a.-p. hippocampus sections of Visium ST; white, nuclei counterstain (DAPI).
(**D**) Spatially resolved detection of 22 abundant, but locally restricted barcodes (blue in Fig. 4E), in 16 a.-p. hippocampus sections of Visium ST; white, nuclei counterstain (DAPI).

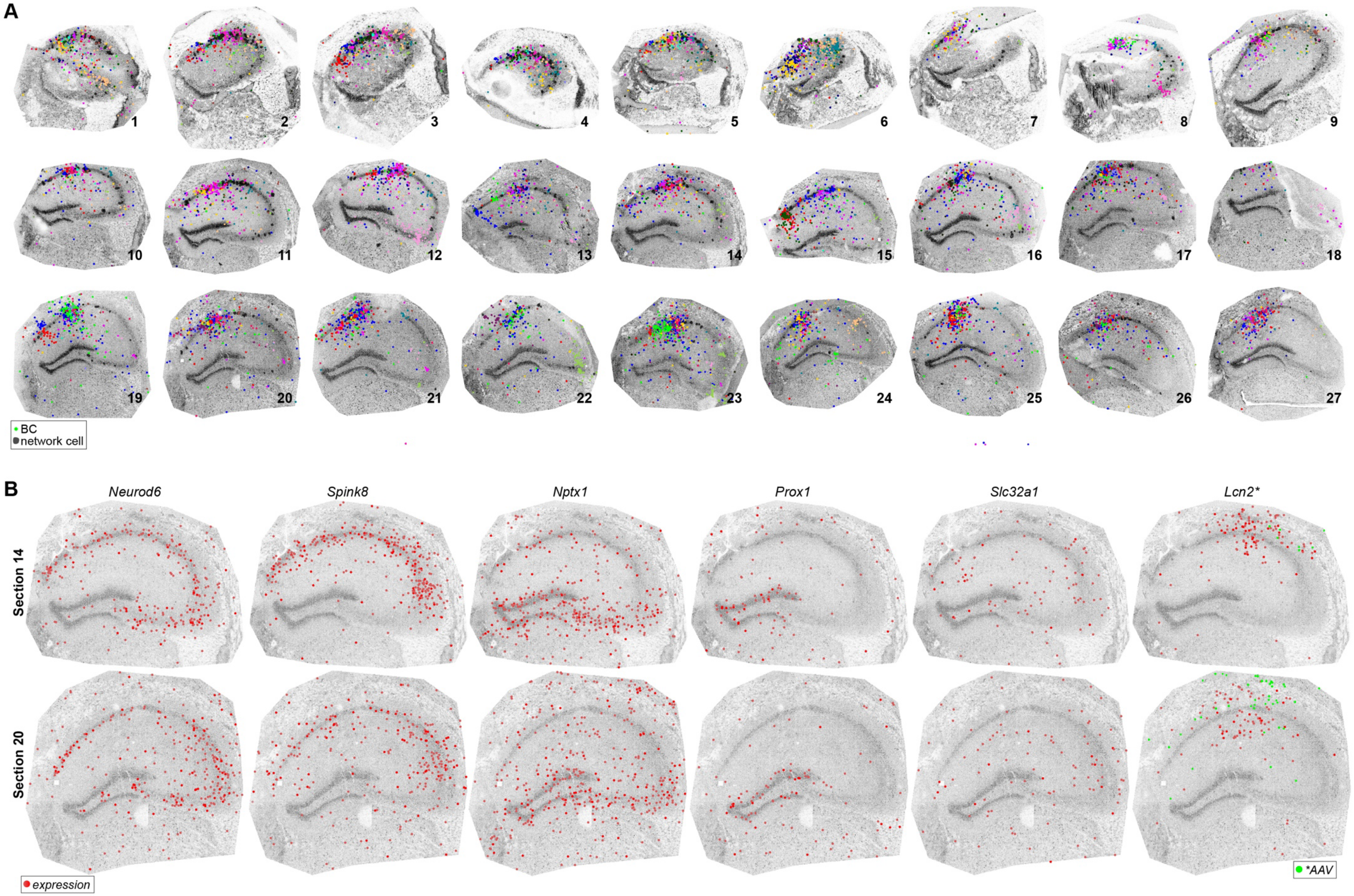
A
BC
network cell
B
Neurod6
Spink8
Nptx1
Prox1
Slc32a1
Lcn2*
Section 14
Section 20
expression
*AAV

**Figure S11 Cell-resolved hippocampus networks on Stereo-seq STOmics**

(**A**) Spatially resolved detection of the top 20 abundant barcodes (colors), in all 27 a.-p. hippocampus sections. Black, network cell outlines. The grey scale showing section anatomy is based on total detected molecules; where dark is the maximum detected molecules.

(**B**) Spatial gene expression of neuronal markers *Neurod6* (pyramidal)*, Spink8* (CA1-CA2)*, Nptx1* (CA3), *Prox1* (DG), *Slc32a1* (inhibitory) and inflammatory marker *Lcn2*, as well as AAV-transcripts (green) to indicate the injection site, in sections 14 and 20. Grey scale, total detected molecules; where dark is maximum.
